# Population genomics supports clonal reproduction and multiple gains and losses of parasitic abilities in the most devastating nematode plant pest

**DOI:** 10.1101/362129

**Authors:** Georgios D. Koutsovoulos, Eder Marques, Marie-Jeanne Arguel, Laurent Duret, Andressa C.Z. Machado, Regina M.D.G. Carneiro, Djampa K. Kozlowski, Marc Bailly-Bechet, Philippe Castagnone-Sereno, Erika V.S. Albuquerque, Etienne G.J. Danchin

## Abstract

The most devastating nematodes to worldwide agriculture are the root-knot nematodes with *Meloidogyne incognita* being the most widely distributed and damaging species. This parasitic and ecological success seem surprising given its supposed obligatory clonal reproduction. Clonal reproduction has been suspected based on cytological observations but, so far, never confirmed by population genomics data. At the species level, *M. incognita* is highly polyphagous with thousands of host plants. However, the host range varies among different *M. incognita* isolates that may present distinct and more restricted host compatibilities. Historically, four ‘host races’ had been defined as a function of ranges of compatible and incompatible plants. We sequenced the genomes of 11 isolates across Brazil, covering these four distinct races to assess (i) how clonal reproduction is and (ii) how the level of genome variability associates with biological traits such as the host races, affected agronomic culture, and geographical distribution. By aligning the genomic reads of the isolates to the *M. incognita* reference genome assembly, we identified SNV and small-scale insertions/deletions. Analysis of linkage disequilibrium and 4-gametes test, showed no sign of recombination, confirming the clonal mode of reproduction of *M. incognita.* We showed that there are relatively few point variations between the different isolates, and these variations show no significant association with either the host races, the geographical origin of the samples or the host plant on which they have been collected. Due to the lack of phylogenetic signal underlying their existence, we recommend the discontinuation of the terminology ‘race’. Overall, these results suggest that multiple gains and losses of parasitic abilities and adaptations to different environmental conditions account for the broad host spectrum and wide geographic distribution of *M. incognita*. Hence, this nematode constitutes a model species to study adaptability without sexual recombination and overall low genomic variations in animals.

## Introduction

Nematodes cause severe damages to the world agricultural production every year with the root-knot nematodes (RKN, genus *Meloidogyne*) being the most economically harmful in all temperate and tropical producing areas [1,2]. Curiously, the most polyphagous RKN species, able to parasitize the vast majority of flowering plants on Earth [3], are described as mitotic parthenogenetic, based on cytogenetics comparisons with outcrossing relatives [4,5]. This suggests absence of meiosis and obligatory asexual reproduction. Among these mitotic parthenogenetic RKN, *M. incognita* is the most widespread and is present, at least, in all the countries where the lowest temperature exceeds 3°C. Greenhouses over the world also extend its geographic distribution [6]. *M. incognita* is so widely distributed that it is not even included on the list of regulated pests [7]. Due to its worldwide distribution and extremely large range of hosts, *M. incognita* has been deemed the most damaging species of crop pest worldwide [3].

However, it has become more and more evident that the whole broad host range of *M. incognita* as well as the other major RKN species is not present in all the individuals within the species but that different ‘populations’ or ‘isolates’ have different and overlapping ranges of compatible hosts [2]. Variations regarding host range within one given species gave rise to the concept of ‘host race’ as soon as 1952 [8]. Although RKN species can be differentiated based on morphological descriptions [9], isozyme phenotypes [10,11], and molecular analysis [12], this is not the case of host races within a species [5]. Consequently, the pattern of compatibility/incompatibility of the nematode interaction with a set of different host plants was standardised into the North Carolina Differential Host Test (NCDHT, [13]) to differentiate races within Meloidogyne spp. In *M. incognita*, all the populations originally tested reproduced on tomato, watermelon, and pepper and none infected peanut, but they differed in their response to tobacco and cotton defining four distinct host races [13] (Table S1). Whether some genetic features are associated with the *M. incognita* races remains unknown. Indeed, diversity studies using RAPD and ISSR markers across eight isolates and the four races of *M. incognita* found no correlation between phylogeny and host races [14–16]. Although in one of these studies [16], two out of three esterase phenotypes were monophyletic in the phylogenetic tree of the *M. incognita* isolates, they did not segregate according to the host races. A different molecular approach to try to differentiate host races was also proposed based on repeated sequence sets in the mitochondrial genome [17]. Although the pattern of repeats allowed differentiating one isolate of race 1, one of race 2 and one of race 4; the study encompassed only one isolate per race, and thus the segregation could be due to differences between isolates unrelated to the host race status itself.

Hence, no clear genetic determinant underlying the phenotypic diversity of *M. incognita* isolates in terms of host compatibility patterns have been identified so far [18]. This lack of phylogenetic signal underlying the host races is surprising because it would suggest multiple independent gains and losses of host compatibly patterns despite clonal reproduction. Theoretically, animal clones have poorer adaptability because the efficiency of selection is impaired, advantageous alleles from different individuals cannot be combined and deleterious mutations are predicted to progressively accumulate in an irreversible ratchet-like mode [19–22].

For these reasons, the parasitic success of *M. incognita* has long been described as a surprising evolutionary paradox [23]. However, this apparent paradox holds true only if this species does actually reproduce without sex and meiosis while presenting substantial adaptability. So far, no whole genome level study conclusively support these tenets.

A first version of the genome of *M. incognita* was initially published in 2008 [24] and re-sequenced at higher resolution in 2017, providing the most complete *M. incognita* reference genome available to date [25]. This study showed that the genome is triploid with high average divergence between the three genome copies most likely because of hybridization events. Due to the high divergence between the homoeologous genome copies, and the supposed lack of meiosis, it was assumed that the genome was effectively haploid. The genome structure itself showed synteny breakpoints between the homoeologous regions and some of them formed tandem repeats and palindromes. These same structures were also described in the genome of the bdelloid rotifer *Adineta vaga* and considered as incompatible with meiosis [25,26]. However, whether these structures represent a biological reality or artefacts of genome assembly remains to be clarified. Indeed, both genomes have been assembled using the same techniques and no independent biological validation for these structures has been performed. Hence, so far no strong evidence supporting the absence of meiosis was available at the genome level.

Furthermore, because the reference genome was obtained from the offspring of one single female (originally from Morelos, Mexico), no information about the genomic variability between different populations or isolates was available. More recently, a comparative genomics analysis, including different strains of *M. incognita*, showed little variation at the protein-coding genome level between strains collected across different geographical locations [27], confirming previous observations with RAPD and ISSR markers [14–16]. However, no attempt was made to associate these few variations with biological traits such as the host-race or geographic origin. Moreover, the variability between isolates at the non-coding level, which represents the majority of the genome, was not described in this initial analysis.

In the present study, we used population genomics analyses to investigate, (i) whether the supposed absence of meiosis is supported by the properties of genome-wide SNV markers between isolates, (ii) the level of variation between isolates at the whole genome level, and (iii) whether these variations follow a phylogenetic signal underlying life history traits such as the host compatibility patterns, the geographic distribution or the current host crop plant.

To address these questions, we have sequenced the genomes of 11 isolates representing the four *M. incognita* host-races from populations parasitizing six crops across geographically different sites in Brazil (Figure 1). We used isozymes profiles, SCAR markers, and the NCDHT to characterize the biological materials before the DNA extraction and high coverage genome sequencing. We identified short-scale variations at the whole genome level by comparing the *M. incognita* isolates to the reference genome (Morelos strain from Mexico). We conducted several SNV-based genetic tests to investigate the evidences (or lack thereof) of recombination. Using two different approaches, we classified the *M. incognita* isolates according to their SNV patterns and investigated whether the classification was associated to the following biological traits: host compatibility, geographical localisation and current host plant.

**Figure 1.**
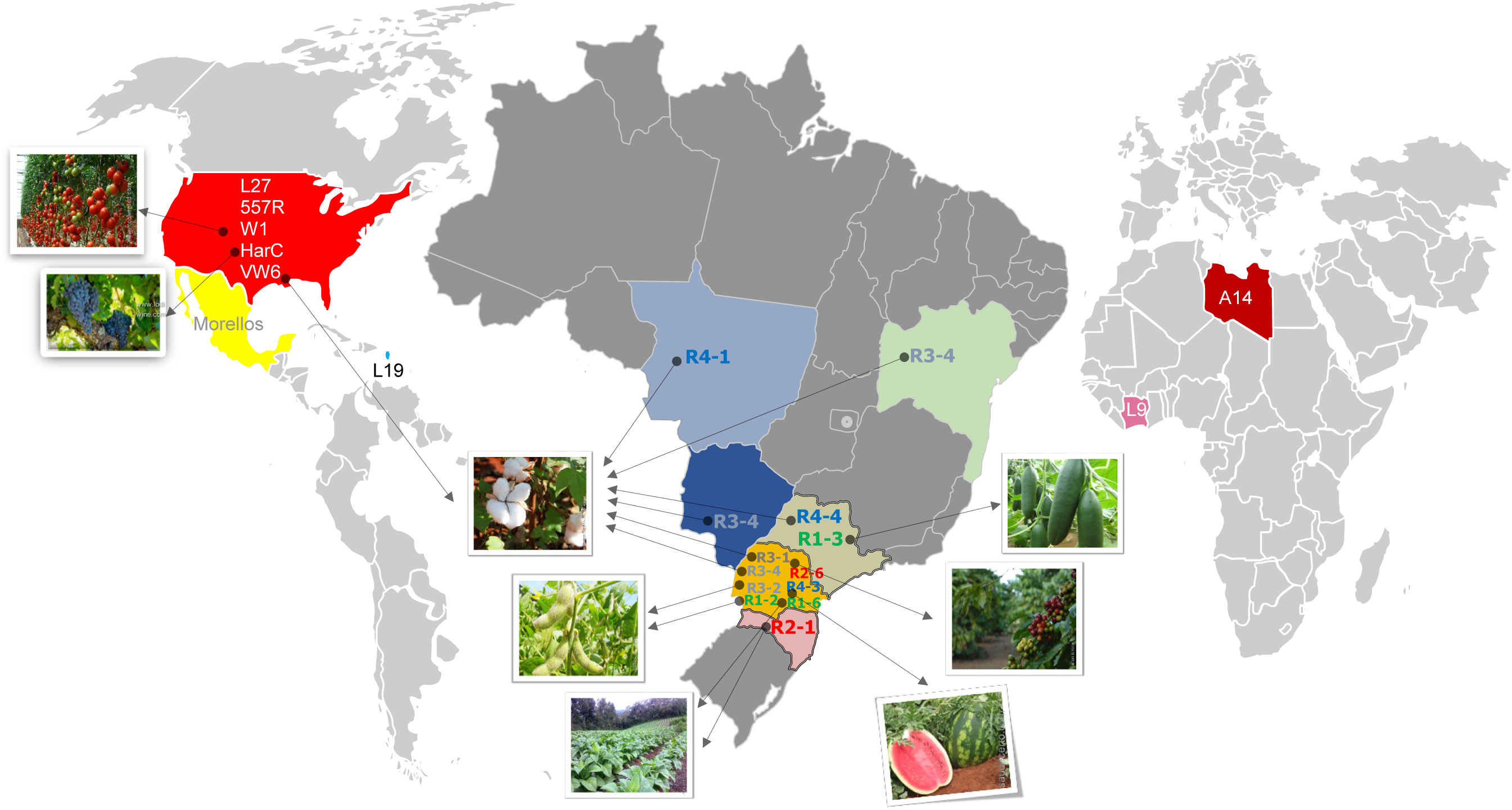
World map showing geographical origins for all samples used in the study. Expanded map of Brazil showing the states where the 11 isolates sequenced in this study were collected. Each state is highlighted with a different colour. The countries listed in the literature for other sequenced genomes are completely coloured. The cultures from which the samples were isolated are illustrated by photographs, which are pointed by arrows coming from the name of the respective isolate. Only 3 of the isolates described previously in the literature have their culture of origin reported. The names of the Brazilian isolates are in 4 different colour sources for each race (race 1 in green, 2 in red, 3 in grey and 4 in blue). The names of the isolates of the literature are written in white or black.

Our population genomics analysis allowed addressing key evolutionary questions such as how asexual is reproduction in this animal species. We were also able to clarify the adaptive potential of this devastating plant pest in relation to its mode of reproduction. In particular, we determined whether there is a phylogenetic signal underlying variations in biological traits of agro-economic importance such as the patterns of host compatibility (host races). While association between phylogenetic signal and patterns of host compatibilities would tend to show stable inheritance from ancestral states, the non-association would support multiple gains and losses of parasitic abilities and substantial adaptability.

This resolution has important agricultural and economic implications since crop rotation and other control strategies should take into account the adaptive potential of this nematode pest.

## Results

### The *M. incognita* genome is mostly haploid and shows few short-scale variations

We collected 11 *M. incognita* populations from six different states across Brazil and from six different crops (soybean, cotton, coffee, cucumber, tobacco, watermelon) (Figure 1). Each isolate was reared by multiplication of the egg mass of one single female on tomato plants (methods). After having confirmed that the 11 isolates we collected showed the characteristic isozyme profiles and molecular signatures of *M. incognita*, we characterized their host race status using the NCDHT (methods, Table 1). We characterised three isolates as race 1, two as race 2, three as race 3, and three as race 4.

**Table 1.**
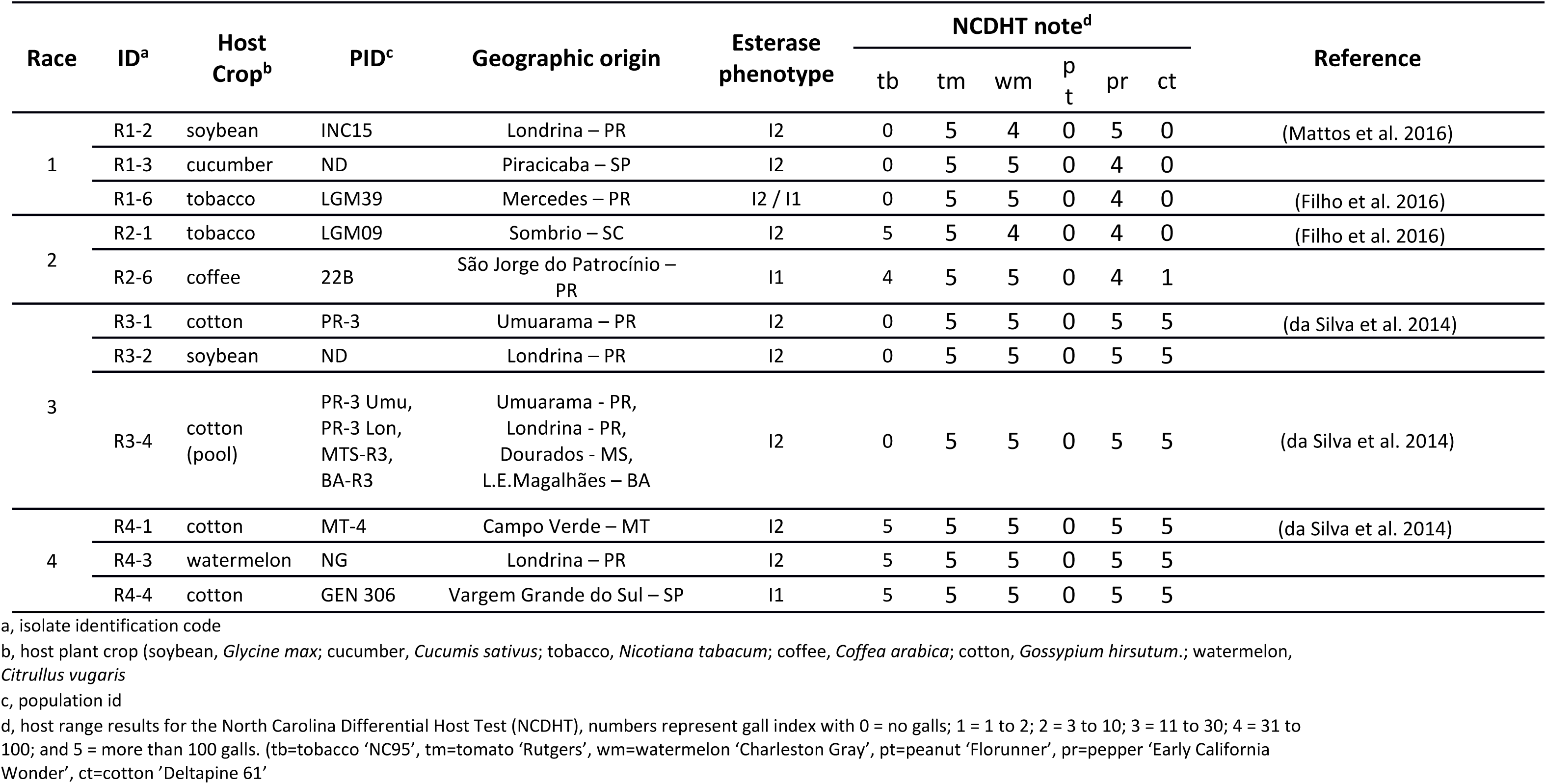
Host race characterization of the 11 *Meloidogyne incognita* isolates used in this study

We generated paired-end genome reads (∼76 million per isolate) which covered the ∼184Mb *M. incognita* genome assembly [25] at a depth > 100X (Table S2) for each isolate. Variant calling, performed in regions with at least 10x coverage per sample, identified 338,960 polymorphic positions (∼0.19% of the total number of non-ambiguous nucleotides). Around 20% of these positions corresponded to 1/1 SNV, fixed within each isolate but variable between isolates and the reference genome. We examined the distribution of base coverage of SNV fixed within all isolates (1/1 fixed SNV) and SNV that presented variations within at least one isolate (0/1 SNV). We observed that the 0/1 SNV, which were variable within isolates, showed a peak of distribution at ∼twice the coverage of the peak for fixed 1/1 SNV in the 11 isolates (Fig. S1). This parallels the distribution of base coverage in the *M. incognita* reference genome scaffolds which shows a major peak at ∼65X and a second minor peak at ∼130X (twice the coverage; Fig. S2). These genome regions at double coverage were considered as representing highly similar pairs of homoeologous genome copies that were collapsed during the assembly [25]. Although these regions are minority in the genome assembly, they seem to be responsible for many 0/1 SNV (presenting within isolate variations). The SNV in these minority regions of double coverage probably result from genome reads of two homoeologous regions mapped to a single collapsed region in the reference assembly. Hence, most of these 0/1 SNV might not represent variations between individuals within an isolate but between the few collapsed homoeologous regions. Because the reference genome is mostly assembled in haploid status (Fig. S2), and the nature of 0/1 SNV is unsure, we will utilise only 1/1 SNV fixed within isolates for all downstream analyses. Although this precludes analyses of variations between individuals within isolates this allows a comparison of variations between isolates based on >66,000 solid fixed markers.

### No evidence for meiotic recombination in *M. incognita*

Based on cytogenetics observation, *M. incognita* and other tropical root-knot nematodes have been described as mitotic parthenogenetic species [4,5]. However, this evolutionary important claim has never been confirmed by genome-wide analyses so far. Using the SNV fixed markers at the whole-genome scale, we conducted linkage disequilibrium (LD) analysis as well as 4-gametes test to search for evidence for recombination (or lack thereof). In an outcrossing species, although physically close markers should be in high LD, this LD should substantially decrease with distance between the markers, because of recombination, and eventually reach absence of LD as for markers present on different chromosomes. In clonal species, however, in the absence of recombination, the LD between markers should remain high and not decrease with distance between markers. By conducting an analysis of LD, we did not find any trend for a decrease of LD between markers as a function of their physical distance (Figure 2). In contrast, the LD values remained high regardless the distance and oscillated between 0.85 – 0.94. Hence, we did not observe the expected characteristic signatures of meiosis. An inversely contrasted situation between outcrossing and clonal genomes should be observed for the 4-gametes test. Taking fixed SNV markers that exist in two states among the 11 isolates, the proportion of pairs of markers that pass the 4-gametes test (i.e. that represent the 4 products of meiosis) should rapidly increase with distance between the markers, in case of recombination. In contrast, in the absence of recombination, no trend for an increase of the proportion of pairs or markers passing the 4-gametes test with distance between markers should be observed. By conducting an analysis of 2-states markers, we observed no trend for a change in the proportion of markers passing the test with distance. In contrast, the distribution remained flat and close to a value of 0.0. Again, this trend does not correspond to the expected characteristic of meiotic recombination.

**Figure 2.**
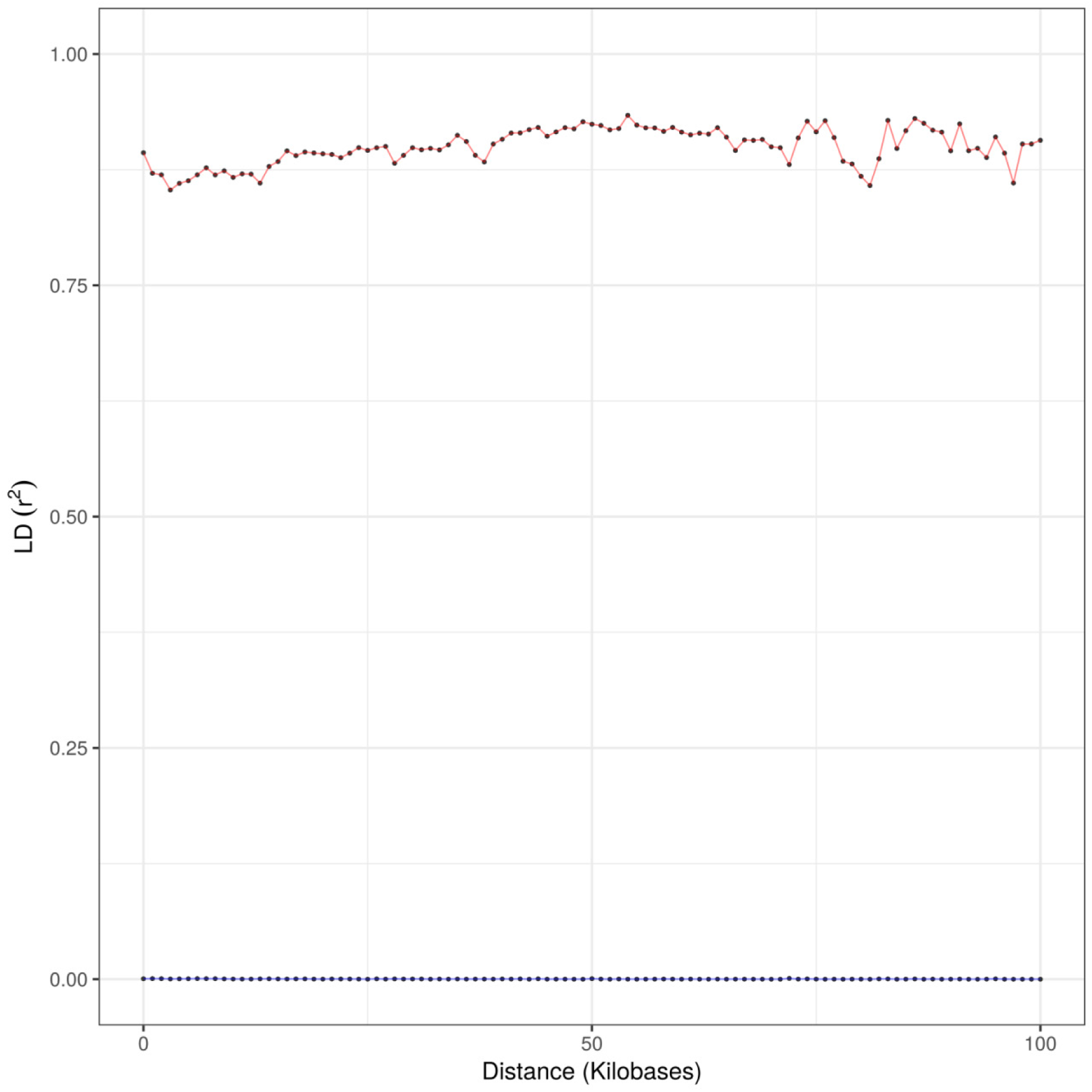
Linkage Disequilibrium and 4-gametes test of *M. incognita* isolates. Based on SNVs fixed within isolates: proportion of pair of markers that pass the 4-gametes test (blue) and linkage disequilibrium measured as r2 between markers (red), both as a function of the inter-markers distance on *M. incognita* scaffolds.

To assess the sensitivity of our method in finding evidence for recombination, we conducted the same analyses (LD and 4-gametes tests) in the outcrossing diploid meiotic plant-parasitic nematode *Globodera rostochiensis* [28]. Because the *G. rostochiensis* genome assembly mostly consists of merged paternal and maternal haplotypes, we had to phase the SNVs before conducting LD and 4-gametes tests. The results were totally contrasted between *M. incognita* and *G. rostochiensis* (Figure 3). In *G. rostochiensis*, the LD and 4-gametes curves started at relatively lower (<0.7) and higher (>0.15) values, respectively. Furthermore, we observed a rapid exponential decrease of r^2^ in the first kb for LD. At an inter-marker distance of 3kb the r^2^ value was <0.37. In parallel, we observed a concomitant rapid and exponential increase in the proportion of markers passing the 4-gametes test, which was >0.38 at the same inter-marker distance. Hence, while *G. rostochiensis* appears to display all the expected characteristics of meiotic recombination, this was not the case for *M. incognita*. This confirms at a whole genome-scale the lack of evidence for meiosis previously observed at the cytological level in *M. incognita*.

**Figure 3.**
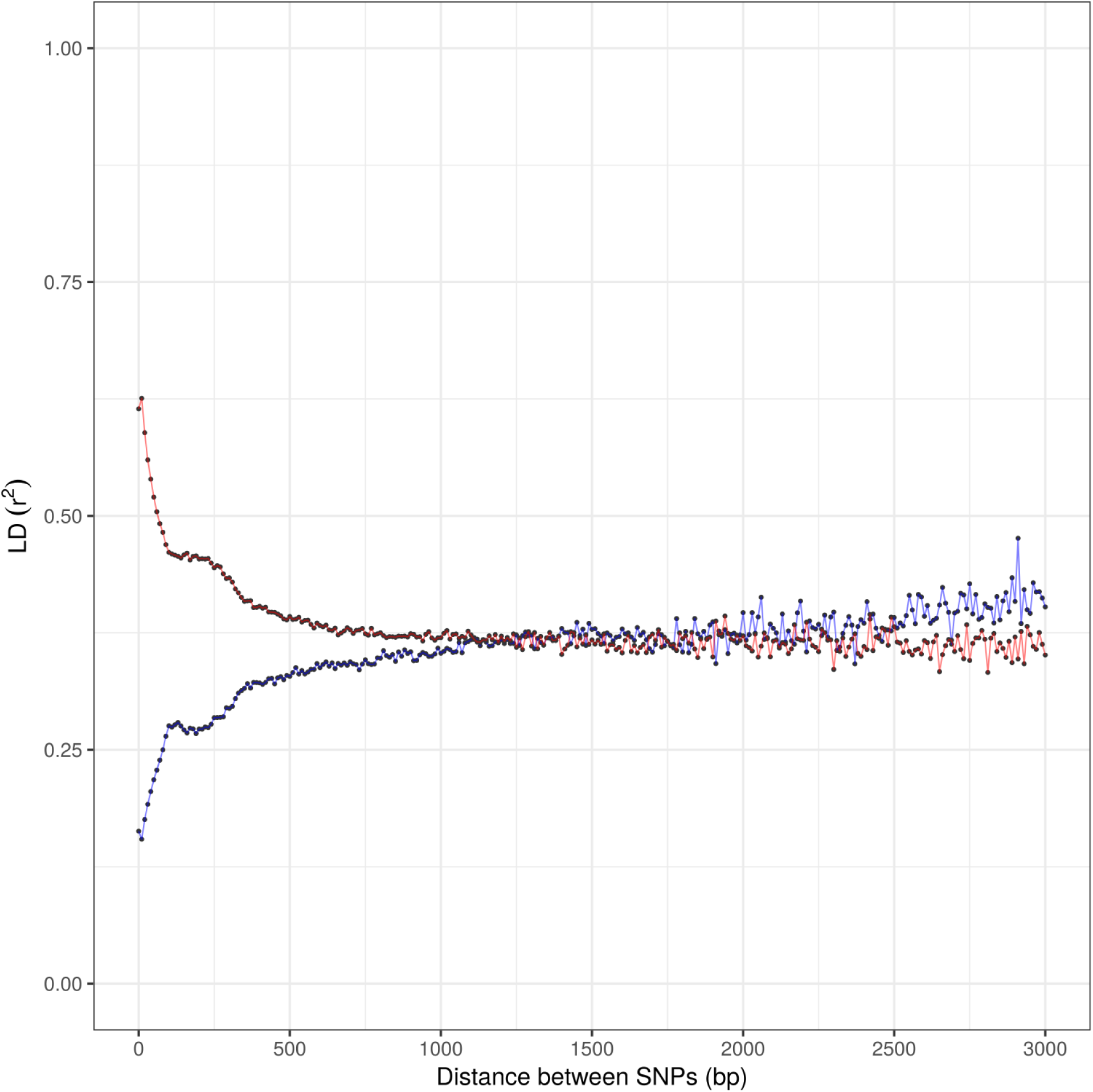
Linkage Disequilibrium and 4 gamete test for phased SNVs in *Globodera rostochiensis* isolates. The r^2^ correlation between markers, indicating linkage disequilibrium (LD) is given in the red (upper) plot, as a function of the physical distance between the markers. The proportion of pairs of two-state markers that pass the 4-gamete test is given in the blue (lower) plot as a function of the distance between the markers.

### The levels of variations between isolates are low and not specific to races

Each isolate showed a different level of divergence from the reference genome with R1-2 having the highest number of fixed SNV (41,518) and R1-6 having the least (17,194) variants (Figure 4). The R3-4 isolate originated from a pool of four populations. However, the low number of SNV compared to the reference indicates either that the genomes of these four populations were very close or that a specific population displaced the other three (Figure 4). Thus, the R3-4 isolate was analysed exactly as the other isolates. Overall, the percentages of fixed SNV on the nuclear genomes of the eleven isolates, compared to the Morelos reference strain, ranged between 0.01 % and 0.02 %. In comparison, the percentages of SNV in the mitochondrial genome ranged between 0.04% and 0.18%.

**Figure 4.**
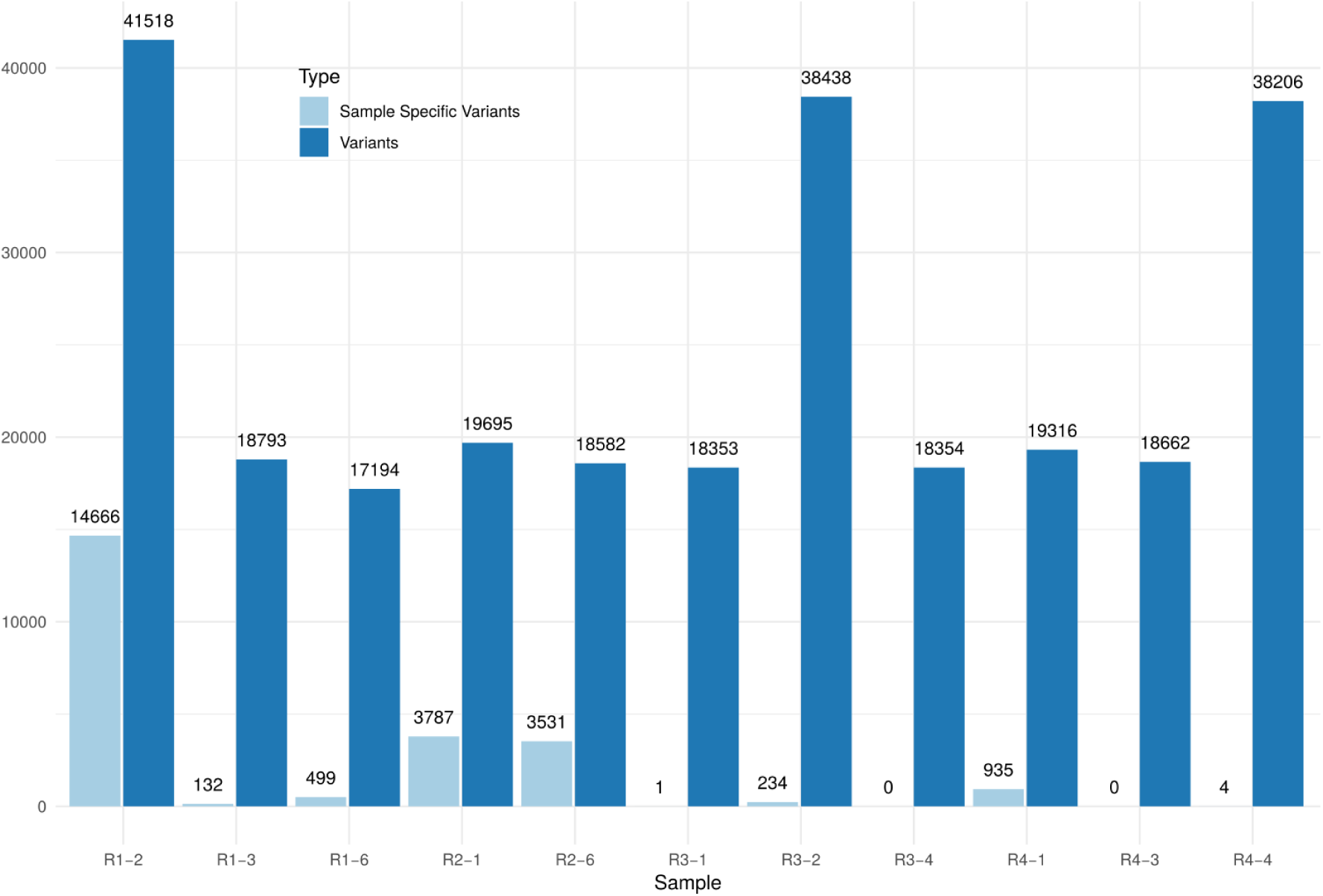
Distribution of the number of variants per race and isolate. Number of variants per isolate and isolate-specific variants for the 11 Brazilian isolates.

Interestingly, race-specific variants exist only for race 2, which exhibited 30 race-specific variations. This is possibly due to the fact that race 2 is represented by only two isolates (vs. 3 for the rest of the isolates). The vast majority (∼78%) of SNV were outside of coding regions; only 14,704 variable positions fell in coding regions and covered 7,259 out of 43,718 predicted protein-coding genes. In these coding regions, 8,179 were synonymous substitutions, 3,854 SNV yielded non-synonymous substitution, 93 nonsense mutations and the rest other disruptive mutations.

From the SNV falling in coding regions, we constructed a multiple alignment and measured nucleotide diversity at synonymous (π_s_) and non-synonymous (π_n_) sites for the 11 isolates as well as the π_n_/π_s_ ratio as a measure of the efficiency of selection. Consistent with the overall low number of SNV, the π_s_ value across the 11 isolates was low (1.29 10^−03^). This is one order of magnitude lower than the values measured for two outcrosser nematodes from the *Caenorhabditis* genus [29], *C. doughertyi* (formerly sp. 10: 4.93 10^−02^) and *C. brenneri* (3.22 10^−02^). A similar difference of one order of magnitude was also observed for the diversity at non-synonymous sites with a π_n_ value of 1.66 10^−04^ for *M. incognita* and values reaching 2.53 10^−03^ and 1.28 10^−03^ for *C. doughetryi* and *C. brenneri*, respectively. However, the π_n_/π_s_ ratio was substantially higher for *M. incognita* (0.129) than for the two outcrossing Caenorhabditis (0.051 and 0.040 for *C. doughetryi* and *C. brenneri*, respectively). These results suggest a lower efficacy of selection in the obligate parthenogenetic *M. incognita* than in these two outcrossing *Caenorhabditis* nematodes.

### There is no significant association between the short-scale variants and biological traits

Using principal component analysis on the whole set of fixed SNV, we showed that the eleven *M. incognita* isolates formed three distinct clusters, which we named A, B and C (Figure 5). Cluster A is represented by isolate R1-2 alone, which has the highest number of variants. Cluster B is constituted by R3-2 and R4-4. The rest of the isolates fall in a single dense cluster C of overall low variation. There was no significant association between the clusters and the host race status (Fisher’s exact test p-value=1, Sup. Text, Table S3). This implies that isolates of the same host race are not more similar to each other than isolates of different host races. There was also no significant association between the SNV-based clusters and the original host plant from which the nematodes have been collected (Fisher’s exact test p-value=0.69, Sup. Text, Table S3). Interestingly, the four different host races are all represented in one single cluster (C). Within this cluster, the total number of variable positions was 29,597. Meaning that the whole range of host-race diversity is present in a cluster that represents only 44% of the total existing genomic variation. We also conducted an isolation by distance (IBD) analysis, which showed no correlation between the genetic distance and the geographical distance (Fig. S3).

**Figure 5.**
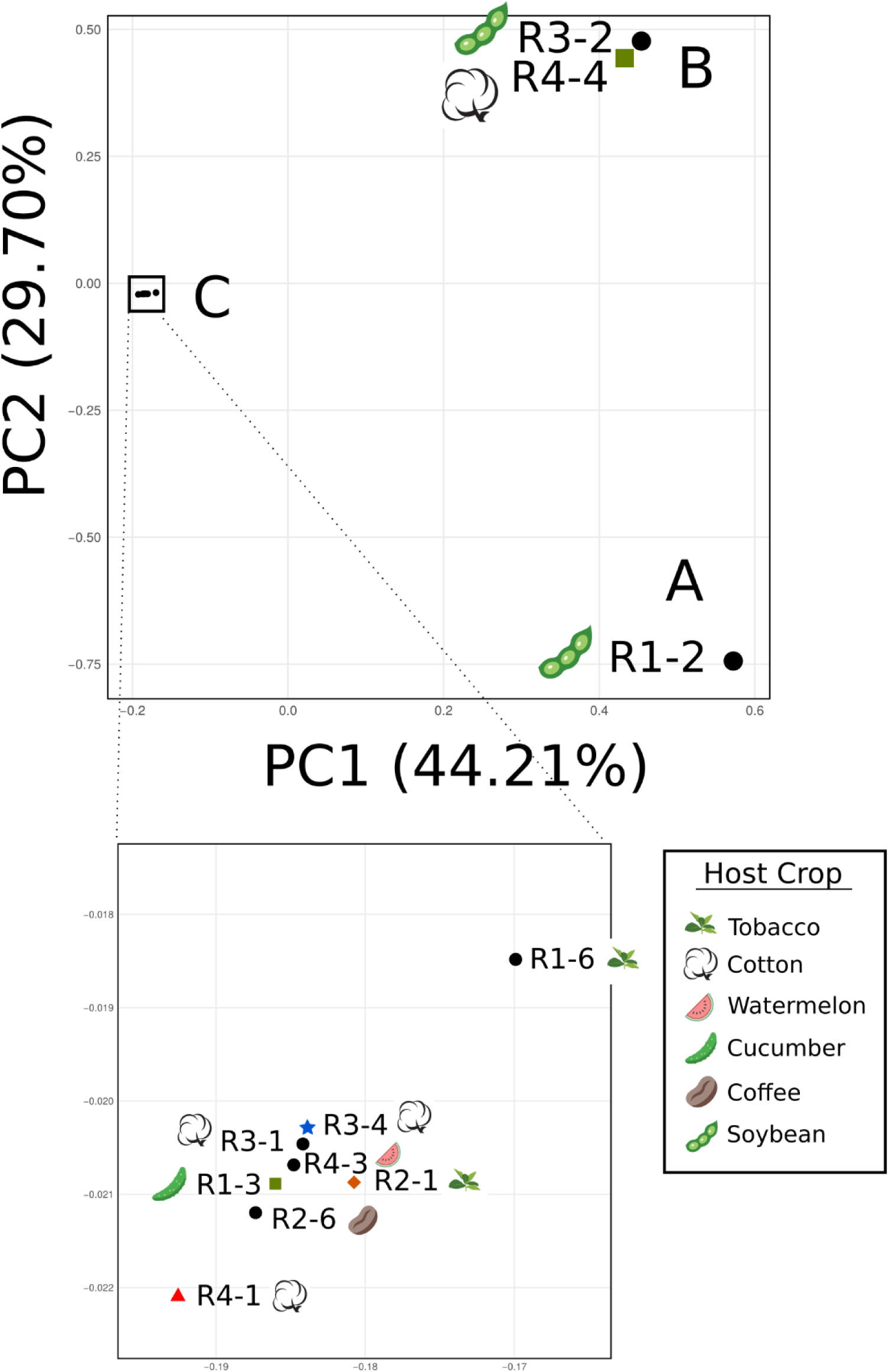
PCA analysis of the different *M. incognita* isolates groups them into three clusters (A, B, and C). The geographic origins are associated to coloured shapes: black circle: Paraná, orange diamond: Santa Catarina, green square: São Paulo, red triangle: Mato Grosso, blue star: pool. Host plant representative pictures are displayed next to the isolates: soybean pod (R1-2 and R3-2); cotton flower (R3-1, R3-4, R4-4, and R4-1); coffee grain (R2-6); cucumber vegetable (R1-3); tobacco leaves (R1-6 and R2-1); and watermelon fruit slice (R4-3).

To assess the levels of separation vs. past genetic exchanges between these clusters, we calculated fixation index values (F_ST_). Weighted F_ST_ values between clusters were all >0.83, suggesting a lack of genetic connections between the clusters (Table S4). Using the mean F_ST_ values, in contrast, while we observed a mean F_ST_ >0.98 between clusters A and B, indicating a lack of genetic connection between R1-2 and cluster B, the F_ST_ values were much lower between A and C (0.35) and between B and C (0.52). This would suggest isolates from clusters A and B both result from a past bottlenecked dispersal and propagation from some isolates in cluster C. We also conducted the same π_n_/π_s_ analysis than the one performed at the whole species level for each cluster of the PCA containing at least 2 isolates. These cluster-specific statistics yielded similar π_n_/π_s_ ratio than the one observed at whole species level (Cluster C: π_s_ 3.8 10^−04^, π_n_ 5.36 10^−05^, π_n_/π_s_ 0.141; Cluster B: π_s_ 2.08 10^−05^, π_n_ 2.64 10^−06^, π_n_/π_s_ 0.127).

### Phylogenetic networks confirm the lack of association of SNV with biological traits and support clonal evolution

Using a phylogenetic network analysis based on SNV present in coding regions, we could confirm the same three clusters (Figure 6). This further supports the absence of phylogenetic signal underlying the host races (patterns of host compatibilities). Interestingly, this network analysis based on fixed SNV yielded a bifurcating tree and not a network. This result suggests a lack of genetic exchanges between the isolates, as expected from a clonal species. To confirm this result, we conducted separate phylogenetic analyses for each of the 14 longest scaffolds with sufficient number of phylogenetically informative variable positions and the mitochondrial genome. All these analyses showed a clear separation between the three clusters (A, B, and C) with some minor polytomies within cluster C (Fig. S5 and Fig; S6).

**Figure 6.**
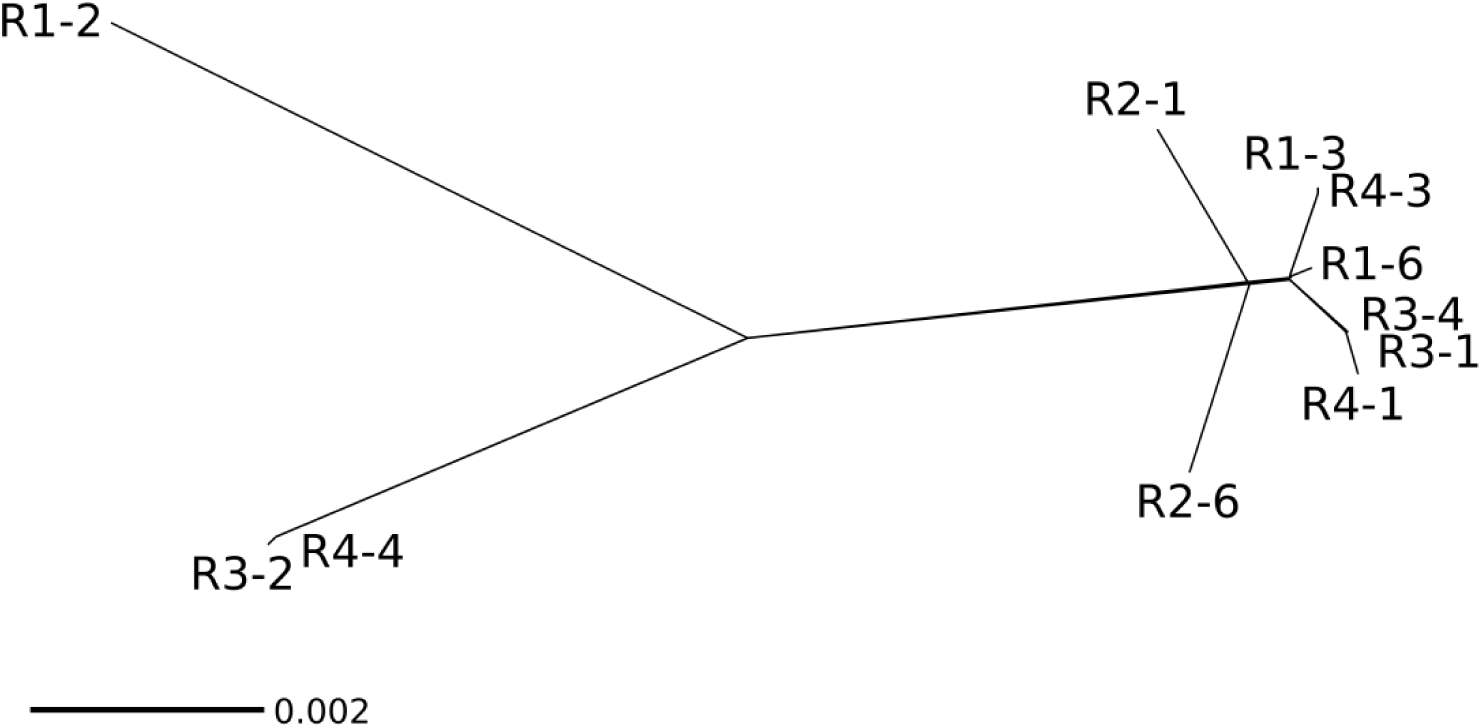
Phylogenetic network for *M. incognita* isolates based on SNV present in coding sequences. The phylogenetic network based only on changes in coding sequences shows the same grouping into 3 distinct groups.

According to the two classification methods (PCA and phylogenetic network), isolate R1-2 seemed to be the most divergent from the rest of isolates, which is consistent with its higher total number of SNV and number of isolate-specific SNV. Then, a small cluster was composed of isolates R3-2 and R4-3 (equivalent to cluster B of the PCA). Finally, a cluster (equivalent to PCA cluster C) grouped the rest of the eight isolates and covered all the defined host races as well as 5 of the 6 different host plants Consistent with the PCA and phylogenetic network analysis, we also did not observe significant association between the number of repeats in the two repeat regions in the mitochondrial genome (63R and 102R) and races, geographical origin or host plant of origin (Table 2).

**Table 2.** Number of repeats per region (63nt and 102nt) in the mitochondrial DNA of each isolate, decimals indicate truncated repeats

| **ID** | **63nt**<br>**Region** | **102nt**<br>**Region** | **Location** | **Host plant** |
| --- | --- | --- | --- | --- |
| **R1-2** | 7.3 | 5.5 | Londrina - PR | soybean |
| **R1-3** | 7 | 13 | Piracicaba - SP | cucumber |
| **R1-6** | 1.2 | 7 | Mercedes - PR | tobacco |
| **R2-1** | 7 | 15.4 | Sombrio - SC | tobacco |
| **R2-6** | 7 | 9 | São Jorge do Patrocínio - PR | coffee |
| **R3-1** | 7 | 13 | Umuarama - PR | cotton |
| **R3-2** | 14 | 8.3 | Londrina - PR | soybean |
| **R3-4** | 6 | 13 | Umuarama – PR<br>Londrina – PR<br>Dourados – MS<br>L.E.Magalhães - BA | cotton |
| **R4-1** | 6 | 14.7 | Campo Verde - MT | cotton |
| **R4-3** | 3 | 9 | Londrina - PR | watermelon |
| **R4-4** | 14 | 8.3 | Vargem Grande do Sul - SP | cotton |

### Addition of further geographical isolates does not increase the genomic diversity and confirms the lack of association between genetic distance and biological traits

To investigate more widely the diversity of *M. incognita* isolates in relation to their mode of reproduction and other biological traits, we included whole-genome sequencing data for additional geographical isolates [27]. These genome data included one isolate from Ivory Coast, one from Libya, one from Guadeloupe (French West Indies) and five from the USA (Figure 1). We pooled these eight new isolates with the eleven Brazilian isolates produced as part of this analysis as well as the *M. incognita* Morelos strain (reference genome) and performed a new PCA with the same methodology. Astonishingly, adding these new isolates recovered the same separation in three distinct clusters (A, B and C) (Figure 7). All the new isolates from additional and more diverse geographical origins fell in just two of the previous Brazilian clusters (A and C). Cluster A that previously contained R2-1 alone, now encompasses the Ivory Coast, Libyan and Guadeloupe isolates. Cluster C that previously contained eight of the Brazilian isolates and covered all the host races now includes the five US isolates as well as the Mexican isolate (Morelos, reference genome). Cluster B remains so far Brazilian-specific with only R3-2 and R4-4 in this cluster. Addition of these new geographical isolates [27] did not substantially increase the number of detected variable positions in the genome. Analyses ran with this whole set of available *M. incognita* isolates also further supported the lack of association of SNV-based clusters with biological traits such as host races, nature of the host of origin and geographical distribution (Text S1, Fig. S7).

**Figure 7.**
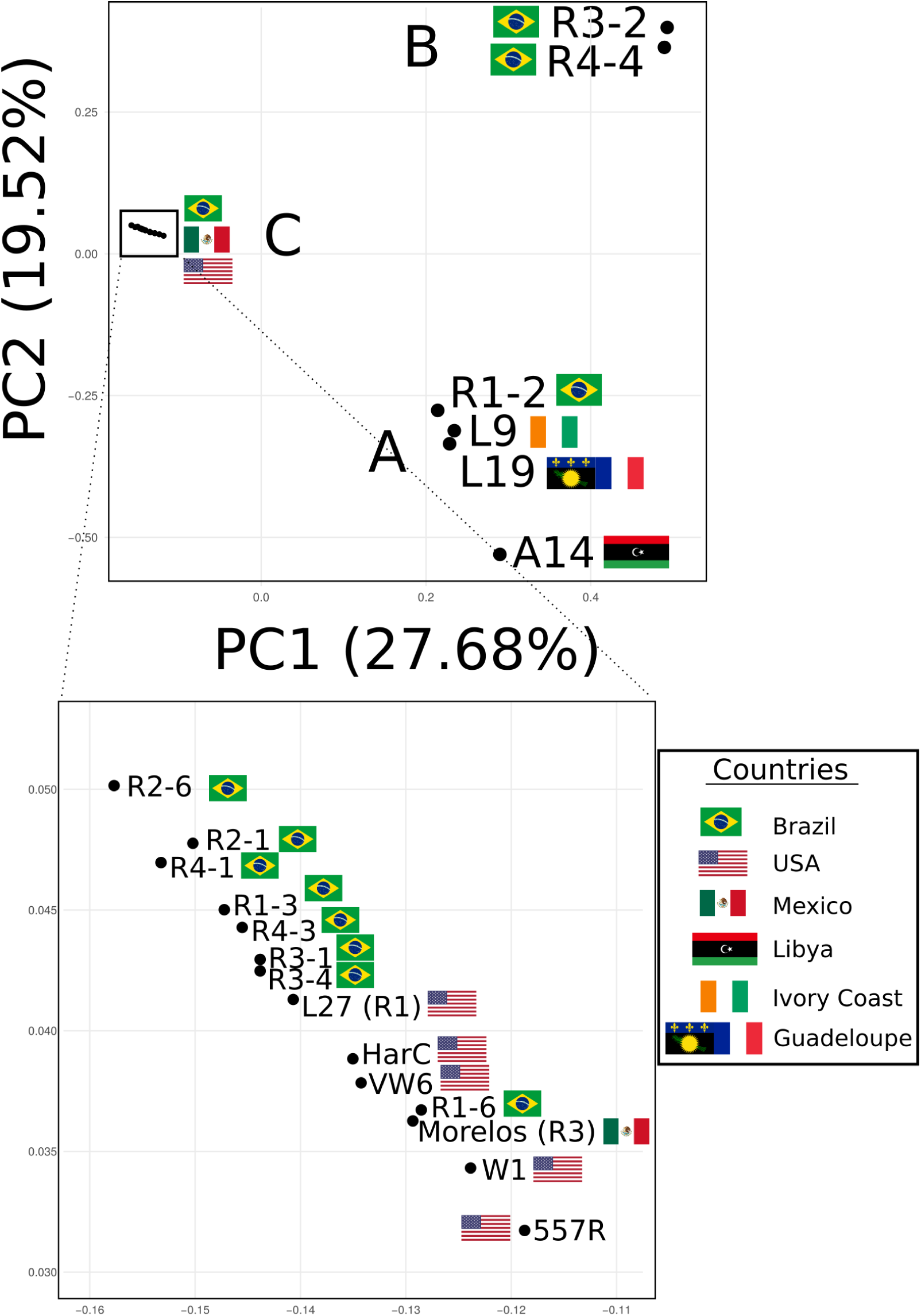
PCA analysis of all known *M. incognita* genomes. The isolates were regrouped based on SNV patterns confirming the same three clusters. Origin countries are indicated by flags (Brazil for R1-2, R1-3, R1-6, R2-1, R2-6, R3-1, R3-2, R3-4, R4-1, R4-3, R4-4; USA for L27, 557R, HarC, W1, VW6; Mexico for Morelos; Libya for A14; Ivory Coast for L9; Guadeloupe for L19).

## Methods

### Purification, species identification of *M. incognita* isolates and determination of host races

The *M. incognita* isolates involved in this study (Table 1) originate from populations collected from different crops and geographically distant origins in Brazil (Figure 1). For each isolate, one single female and its associated egg mass were retrieved as explained in [30]. To determine the species (here *M. incognita*), we used esterase isozyme patterns on the female [10]. The corresponding single egg mass was used for tomato plant infection and multiplication. We reproduced egg-mass isolates on tomato plants (*Solanum lycopersicum* L. cv. Santa Clara) under greenhouse conditions at a temperature of 25-28 ° C. After three months, we confirmed the *M. incognita* species using esterase phenotypes [30]. Once enough nematodes were multiplied, a pool was collected and we performed the North Carolina Differential Host Test (NCDHT) [13] with the following plants: cotton cv. Deltapine 61, tobacco cv. NC95, pepper cv. Early California Wonder, watermelon cv. Charleston Gray, peanut cv. Florunner and tomato cv. Rutgers to determine the host race status. We inoculated these plants with 5,000 eggs and J2 of *M. incognita* and maintained them under glasshouse conditions at 25-28° C for three months, with watering and fertilisation as needed. Two months after inoculation, the root system was rinsed with tap water, and egg masses were stained with Phloxine B [13] to count the number of galls and eggs masses separated for each root system. We assigned a rating index number according to the scale: 0 = no galls or egg masses; 1 = 1-2 galls or egg masses; 2 = 3-10 galls or egg masses; 3 = 11-30 galls or egg masses; 4 = 31-100 galls or egg masses; and 5 > 100 galls or egg masses per root system (Table 1). Host–plants types that have an average gall and egg mass index of 2 or less are designated non-host (-). The other plants (index ≥ 3) are designated hosts (+). We categorised *M. incognita* host races based on their ability to parasitize tobacco and cotton (Table 1). Classically, the index for Rutgers tomato (susceptible control) is higher than 4 (+) [13].

The rest of the population was kept for multiplication on tomato plants to produce enough nematodes for sequencing (typically >1 million individuals pooled together).

### DNA preparation and SCAR test

For each characterized nematode isolate, we extracted and purified the genomic DNA from pooled eggs with the supplement protocol for nematodes of the QIAGEN Gentra^®^ Puregene^®^ Tissue Kit with the following modifications: incubation at 65 ° C in the cell lysis buffer for 30 min and incubation at 55 ° C with proteinase K for 4h. We verified DNA integrity on 0.8% agarose gel and the DNA quantification on Nanodrop. We confirmed isolate species purity by SCAR-PCR [31,32] using the SCAR primers specified in Table S5 for the RKN *M. javanica, M. paranaensis, M. incognita*, and *M. exigua*.

### Sequencing library preparation

We assessed input gDNA quantity using Qubit and normalised the samples to 20ng/ul as described in TruSeq^®^DNA PCR-Free Library Prep Reference Guide (#FC-121-3001, Illumina) prior fragmentation to 350bp with Covaris S2. We assessed the quality of fragments after size selection and size control of the final libraries using High Sensitivity DNA Labchip kit on an Agilent 2100 Bioanalyzer.

### Whole genome sequencing

We quantified isolated sample libraries with KAPA library quantification kit (#7960298001, Roche) twice with two independent dilutions at 1:10,000 and 1:20,000. We calculated the average concentration of undiluted libraries and normalised them to 2nM each then pooled them for sequencing step.

We generated high-coverage genomic data for the 11 *M. incognita* isolates by 2×150 bp paired-end Illumina NextSeq 500 sequencing with High Output Flow Cell Cartridge V2 (#15065973, Illumina) and High Output Reagent Cartridge V2 300 cycles (#15057929, Illumina) on the UCA Genomix sequencing platform, in Sophia-Antipolis, France. We performed two runs to balance the read’s representation among the isolates and obtain homogeneity of coverage for the different samples (Table S2).

### Variant Detection

We trimmed and cleaned the reads from each library with cutadapt tool [33] to remove adapter sequences and bases of a quality inferior to 20. We mapped the clean reads to the *M. incognita* reference genome [25], using the BWA-MEM software package [34]. This reference genome is described as triploid with three equally highly diverged A, B and C genome copies as a result of hybridization events. Most of the triplicated regions have been correctly separated during genome assembly, according to genome assembly size (183.53 Mb) that is in the range of the estimated total DNA content in cells via flow cytometry (189±15Mb) [25]). Hence, the genome was considered in this analysis as mostly haploid. However, the distribution of per-base coverage on the genome assembly presented a 2-peaks distribution with a second minor peak at ∼twice the coverage of the main peak (Fig. S2). Genome regions of double coverage most likely represent cases where two of the three homoeologous loci have been collapsed during the assembly, probably due to lower divergence. Such regions will systematically be responsible for ‘artefactual’ 0/1 SNV (presenting variations within isolates) as the reads from the two homoeologous copies will map a single collapsed region in the reference genomes. To avoid confusion between SNV representing true variations between individuals within isolates from those being artefacts due to collapsed homoeologous regions, 0/1 SNV were discarded from the analysis and only 1/1 SNV fixed within isolates were considered.

We used SAMtools [35] to filter alignments with MAPQ lower than 20, sort the alignment file by reference position, and remove multi-mapped alignments.

We used the FreeBayes variant detection tool [36] to call SNV and small-scale insertions/deletions, incorporating all the library alignment files simultaneously and produced a variant call file (VCF). We filtered the resulting VCF file with the vcffilter function of vcflib [37], retaining the positions that had more than 20 Phred-scaled probability (QUAL) and a coverage depth (DP) > 10. To conduct the same analyses on the genome of the meiotic diploid nematode *Globodera pallida*, we first phased the SNV to haplotypes using WhatsHap [38] because the genome assembly mainly consist of collapsed paternal and maternal haplotypes.

### Genetic tests for detection of recombination

We used custom made scripts (cf. Data Accessibility section) to calculate the proportion of fixed markers passing the 4-gametes test and Linkage Disequilibrium (LD) r^2^ values as a function of inter-marker distance along the *M. incognita* genome scaffolds.

### Genetic diversity between isolates, clusters and efficacy of purifying selection

We used bpppopstats from the Bio++ libraries [39] to estimate the nucleotide variability at non-synonymous and synonymous sites as well as efficacy of purifying selection (π_N_, π_S_ and π_N_/π_S_) using a multiple alignment of the coding regions. We calculated fixation index (F_ST_) for the three clusters using vcftools [40].

### Principal component analysis

We performed a principal component analysis (PCA) to classify the isolates according to their SNV patterns and mapped the race characteristics, geographic location, or current host plants on this classification. We used the filtered VCF file as input in the statistical package SNPRelate [41] to perform the PCA with default parameters.

### Phylogenetic analysis

Based on the VCF file and the *M. incognita* gene predictions [25], we selected 85,413 positions that contained synonymous or non-synonymous mutations (i.e. in coding regions). We aligned these positions and then used them as an input in SplitsTree4 with default parameters. The resulting network produced a bifurcating tree that was identical to the one obtained with RAxML-NG using GTR+G+ASC_LEWIS model. The bifurcating tree was used as input to PastML [42] for reconstruction of the ancestral states of ability to parasitize tobacco and cotton (Fig. S8). Phylogenetic inferences for the largest scaffolds containing at least 20 SNV and the mitochondrial genome were conducted with RAxML-NG [43] using the GTR+G substitution model (except for scaffolds 10 and 20 for which the K80+G model was used because not enough phylogenetically informative positions were available).

### Test for association between biological traits and genetic clusters

We used a Fisher’s exact test in R to assess whether there was a significant association between the SNV-based clusters and the host races or the crop species from which the isolates were originally collected. We also conducted an Isolation By Distance (IBD) analysis using the adegenet R package [44] to check how well the genetic distances correlate with geographic distances between the sampling points of the isolates. Geographic distances were calculated from exact sampling locations, when available, or centre points if the region was known but not the exact sampling location. Sample R3-4 was excluded from this analysis since it was a mix of samples pooled together from different geographical locations. L27 was also excluded since the sampling location was unknown.

- Mitochondrial genome analysis

We sub-sampled genomic clean reads to 1% of the total library for each *M. incognita* isolate. Then, we assembled them independently using the plasmidSPAdes assembly pipeline [45]. We extracted the mitochondrial contigs based on similarity to the *M. incognita* reference mitochondrial genome sequence (NCBI ID: NC_024097). In all cases, the mitochondrion was assembled in one single contig. We identified the two repeated regions (63 bp repeat and 102 bp repeat), described in [17] and we calculated the number of each repeat present in these regions.

## Discussion

Is the parasitic success of *M. incognita* an evolutionary paradox? This proposition would be true only if *M. incognita* is adaptive despite having a fully parthenogenetic reproduction. Our results support these two aspects.

The lack of sexual reproduction in *M. incognita* was so far only assumed based upon initial cytogenetic observations [4,5] but never further supported at whole-genome scale. Here, the different analyses we performed at the population genomics level converge in supporting the lack of recombination and genetic exchanges in *M. incognita*. The phylogenetic network analysis based on fixed SNVs returned a bifurcating tree that separated the different isolates and not a network. This suggests a lack of genetic exchange between the isolates. In sexual ‘recombining’ species, the mitochondrial genome accumulates mutations much faster than the nuclear genome. This is also true in the model nematode *C. elegans* where the mitochondrial mutation rate is at least two orders of magnitude higher than the nuclear mutation rate [46,47]. The higher mitochondrial accumulation of mutations is supposed to be the combined result of extremely rare or total lack of recombination, the low effective population size and the effectively haploid inheritance in mitochondria [48]. In *M. incognita*, as opposed to *C. elegans*, we found that the percentage of variable positions in the mitochondrial genome is only one order of magnitude higher than in the nuclear genome. This suggests that the nuclear genome evolves at a comparable rate to the mitochondrial genome and reinforces the idea that the nuclear genome is mostly effectively haploid and non-recombining. Theoretically, the efficacy of selection should be lower in non-recombining species than recombining ones. We showed that the ratio of diversity at non-synonymous sites / diversity at synonymous sites (π_n_/π_s_) was indeed one order of magnitude higher in *M. incognita* than in two outcrossing Caenorhabditis species. Finally, the proportion of markers passing the 4-gametes test and linkage disequilibrium did not show the exponential increase, respectively decrease, with physical distance as expected under recombination. In contrast, a rapid exponential decrease of linkage disequilibrium was recently observed and considered as an evidence for recombination in the bdelloid rotifer *Adineta vaga* [49]. Collectively, these results strongly suggest absence (or extremely rare) recombination and support the mitotic parthenogenetic reproduction of *M. incognita*.

Despite its clonal reproduction, it was already evident that *M. incognita* has an adaptive potential. Indeed, experimental evolution assays have shown the ability of *M. incognita* to overcome resistance conferred by the Mi gene in tomato in a few generations [18,50]. Naturally virulent *M. incognita* populations (i.e. not controlled by the resistance gene) have also been observed in the fields and probably emerged from originally avirulent populations [51–53], although it is unknown if this resistance breaking is as rapid as under controlled lab conditions. However, adapting from a compatible host plant to another very different incompatible plant is certainly more challenging than breaking down a resistance gene in a same plant. Here, we showed that the different host races defined in *M. incognita* as a function of patterns of (in)compatibilities with different plants do not follow a phylogenetic signal. This would imply multiple independent gains and losses of parasitic abilities to arrive at the current phylogenetic distribution of host compatibility patterns (i.e. host races). Whether these multiple gains and losses occurred from a hyper-polyphagous common ancestor or an ancestor with a more restricted host range remains to be clarified. To address this question, we have reconstructed host compatibilities at each ancestral node based on the SNP-based phylogenetic classification of the *M. incognita* isolates (Fig. S8). This reconstruction showed that the two hypotheses concerning the host range status of the last common ancestor were equally likely. Addition of other isolates characterized for their host race might allow to favour one or the other hypothesis in the future. Consistent with multiple gains and losses of parasitic abilities, host race switching within an isolate over time has already been observed. Isolates of *M. incognita* race 2 and 3, which parasitize tobacco and cotton plants respectively, switched to behaviour similar to race 3 and 2 after staying for eight months on coffee plants (Rui Gomes Carneiro, personal communication). Together with the previously reported ability to break down resistance gene in plants, the ability of *M. incognita* to loose and gain ability to infect different plants highlights its adaptive potential.

Overall, we provided here additional evidence for adaptability and the first whole-genome level assessment for the lack of recombination in *M. incognita*, consolidating this species as a main model to study the paradox of adaptability and parasitic success in the absence of sexual reproduction.

The adaptability of *M. incognita* despite its obligatory asexual reproduction and the lack of phylogenetic signal underlying the host races have important practical implications at the agricultural level. Characterizing populations that differ in their ability to infest a particular host (that carries specific resistance genes) is of crucial importance for growers and agronomists. Indeed, the main *Meloidogyne* spp. control strategies consist in deploying resistant cultivars and appropriate crop rotation against a specific given race. If the identity of a population is unknown, the crop selected for use in a management scheme may cause dramatic increases in nematode populations [13]. However, the adaptability of *M. incognita* casts serious doubts on the durability of such strategies and must be taken into account in rotation schemes. Furthermore, the biological reality of host races themselves is challenged by the lack of underlying genetic signal. Actually, the initial host race concept, has never been universally accepted, in part because it covered only a small portion of the whole potential variation in parasitic ability [2]. Although *M. incognita* was already known to parasitize hundreds of host plants, only six different host standards were used to characterise four races. New host races might be defined in the future when including additional hosts in the differential test. Furthermore, using the same six initial host plant species, two additional *M. incognita* races that did not fit into the previously published race scheme have already been described [54]. Although the terminology ‘races’ of *Meloidogyne* spp. has been recommended not to be used since 2009 [2], several papers related to *M. incognita* diversity of host compatibility or selection of resistant cultivars are still using this term; including on coffee [55,56]; cotton [57,58] or soybean [59]. This reflects the practical importance to differentiate *M. incognita* populations according to their different ranges of host compatibilities. However, because these variations in host ranges are not monophyletic and thus do not follow shared common genetic ancestry, we recommend abandoning the term ‘race’.

Another main question relates to the level of intra-specific genome polymorphism required to cover the different ranges of host compatibilities in *M. incognita* and their ability to survive in different environments, despite their clonal reproduction. In this study, we found that the cumulative fixed divergence across the eleven isolates from Brazil and the reference genome (sampled initially from Mexico) reached ∼0.02% of the nucleotides. Addition of isolates from Africa, the West Indies and the USA did not increase the maximal divergence. This relatively low divergence is rather surprising, considering the variability in terms of distinct compatible host spectra (host races). Host-specific SNV were found only for Race 2 and no functional consequence for these SNPs could be found, as they did not fall in predicted coding or evident regulatory regions. Furthermore, the existence of race-specific SNV themselves is even questionable as addition of other isolates might disqualify the few Race 2-specific SNPs in the future. Similarly, when grouped by host plant species there were no disruptive variations identified in the coding regions, we found no SNV associated to cotton, only one synonymous variant for soybean, and only one synonymous variant for tobacco.

Collectively, our observations indicate that *M. incognita* is versatile and adaptive despite its clonal mode of reproduction. The relatively low divergence at the SNV level suggests acquisition of point and short scale mutations followed by selection of the fittest haplotype is probably not the main or at least not the sole player in the adaptation of this species to different host plants and environments. Other mechanisms such as epigenetics, copy number variations, movement of transposable elements or large-scale structural variations could be at play in the surprising adaptability of this clonal species. Consistent with this idea, convergent gene copy number variations (CNV), have recently been shown to be associated with adaptation to a resistance gene-bearing plant in *M. incognita* [60]. Interestingly, the parthenogenetic marbled crayfish has multiplied by more than 100 its original area of repartition across Madagascar, adapting to different environments despite showing a surprisingly low number of nucleotide variation (only ∼400 SNV on a ∼3Gb genome representing a proportion of variable positions of 1.3 10^−7^ only). This also led the authors to suggest that mechanisms other than acquisition of point mutations and selection of the fittest haplotype must be involved [61].

Previously, we have shown that the genome structure of *M. incognita* itself, could participate in its versatility. Indeed, being allopolyploid, *M. incognita* has >90% of its genes in multiple copies. The majority of these gene copies show diverged expression patterns one from the other and signs of positive selection between the gene copies have been identified [25]. How the expression patterns of these gene copies vary across different geographical isolates with different host compatibilities would be interesting to explore in the future.

## Data accessibility

All the sequence data generated during this study have been deposited at the NCBI under GEO accession GSE116847 and available at this URL: https://www.ncbi.nlm.nih.gov/geo/query/acc.cgi?acc=GSE116847 The different scripts and R codes used to process the data are available on GitHub at the following URL:https://github.com/GDKO/gdk_scripts/tree/master/popgenvcf

## Supplementary material

Supplementary figures, text and tables are available online in bioRxiv.

## Acknowledgements

This work was supported by the Agence Nationale de la Recherche program (grant number ANR-13-JSV7-0006—ASEXEVOL), the Insitut National de la Recherche Agronomique program (grant number Accueil de Chercheur Etranger SPE 2016). GK has received the support of the EU in the framework of the Marie-Curie FP7 COFUND People Programme, through the award of an AgreenSkills+ fellowship (under grant number 609398), EM received a post-doc fellowship from the Consórcio Pesquisa Café, EVSA received funding from Embrapa for sequencing at UCA Genomix platform. We thank the SPIBOC Bioinformatics platform for the data hosting.

This preprint has been peer-reviewed and recommended by Peer Community In Evolutionary Biology (https://doi.org/10.24072/pci.evolbiol.100077)

## Author contributions

EGJD, PC-S, GDK, EVSA and LD contributed to the design of the project; RMDGC, ACZM, EM contributed to collection of samples. Classification of samples into races was done by RMDGC, ACZM, EM; library preparations were performed by M-JA; GDK, EGJD, LD, EVSA, MBB and DKK contributed to the analysis and interpretation of data. EGJD and GDK wrote the manuscript with contributions of EVSA, RMDGC, PC-S, M-JA and LD. All authors approved the final manuscript.

