## Supplementary Material for "Population genomics supports clonal reproduction and multiple gains and losses of parasitic abilities in the most devastating nematode plant pest"

| Species | Tobacco<br>*NC 95* | Tomato<br>*Rutgers* | Watermelon<br>*Charleston Gray* | Peanut<br>*Florunner* | Pepper<br>*Early California Wonder* | Cotton<br>*Deltapine 61* |
|---|---|---|---|---|---|---|
| ***M. incognita*** | | | | | | |
| Race 1 | - | + | + | - | + | - |
| Race 2 | + | + | + | - | + | - |
| Race 3 | - | + | + | - | + | + |
| Race 4 | + | + | + | - | + | + |
| ***M. arenaria*** | | | | | | |
| Race 1 | + | + | + | + | + | - |
| Race 2 | + | + | + | - | - | - |
| ***M. javanica*** | + | + | + | - | - | - |
| ***M. hapla*** | + | + | - | + | + | - |

Table S1. Expected host reactions of the NCDHT to discriminate *M. incognita* races.

Nematode reproduction indexes (average of 4 replicates) were used to determine cotton and tobacco as good/susceptible (+) or poor/resistant (-) hosts. Tobacco and cotton are highlighted (light grey shadow) to show that the differentiation of the four *M. incognita* races is determined by the combined reaction of these two hosts alone.

| | Raw | | After Trimming | | | |
|---|---|---|---|---|---|---|
| Isolate | PE reads # | No. of bases | PE reads # | No. of bases | Coverage | Mapped (%) |
| R1-2 | 77.0E+06 | 23.1E+09 | 76.4E+06 | 22.7E+09 | 122.53 | 97.39 |
| R1-3 | 76.2E+06 | 22.9E+09 | 75.5E+06 | 22.4E+09 | 121.20 | 99.51 |
| R1-6 | 75.8E+06 | 22.8E+09 | 75.0E+06 | 22.3E+09 | 120.55 | 97.28 |
| R2-1 | 75.7E+06 | 22.7E+09 | 75.1E+06 | 22.3E+09 | 120.41 | 98.96 |
| R2-6 | 76.0E+06 | 22.8E+09 | 75.3E+06 | 22.4E+09 | 121.01 | 94.26 |
| R3-1 | 75.1E+06 | 22.5E+09 | 74.5E+06 | 22.1E+09 | 119.51 | 89.34 |
| R3-2 | 75.3E+06 | 22.6E+09 | 74.7E+06 | 22.2E+09 | 119.92 | 98.16 |
| R3-4 | 75.3E+06 | 22.6E+09 | 74.6E+06 | 22.2E+09 | 119.79 | 98.47 |
| R4-1 | 75.6E+06 | 22.7E+09 | 75.1E+06 | 22.3E+09 | 120.52 | 99.04 |
| R4-3 | 75.8E+06 | 22.8E+09 | 75.2E+06 | 22.4E+09 | 120.89 | 97.00 |
| R4-4 | 75.6E+06 | 22.7E+09 | 75.0E+06 | 22.3E+09 | 120.46 | 99.32 |

Table S2. Statistics of generated sequence data and genome coverage per isolate.

Number of Illumina paired-end reads and amount of bases generated per isolate. Values indicated before and after quality filtering (trimming). Coverage values of mapped reads are given according to a reference genome size of 184 Mb.

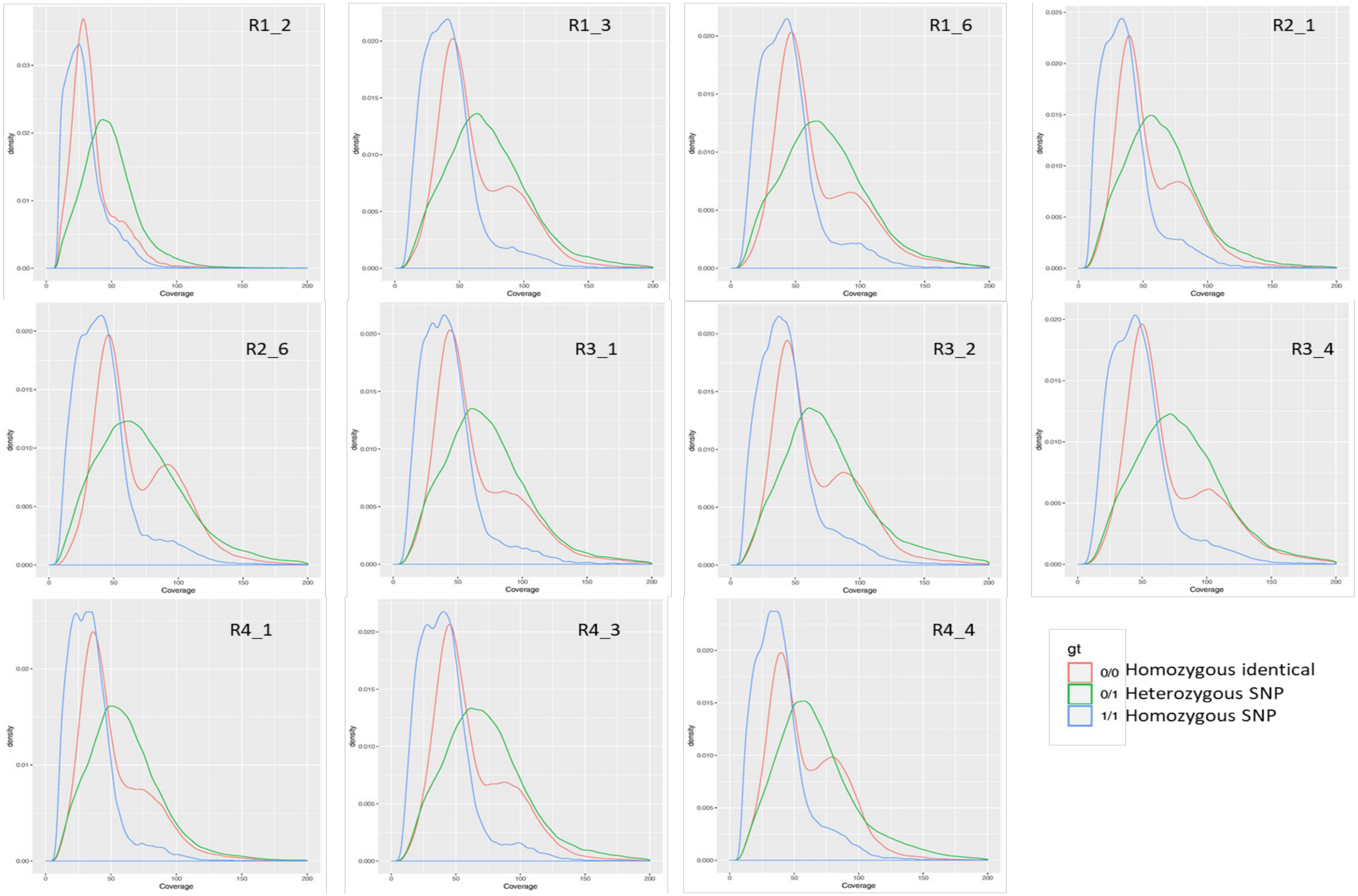


Fig. S1. Distribution of base coverage per category of SNP on the reference genome for each isolate.

Density plot of base coverage for homozygous identical (0/0) heterozygous SNP (0/1) and homozygous SNP.

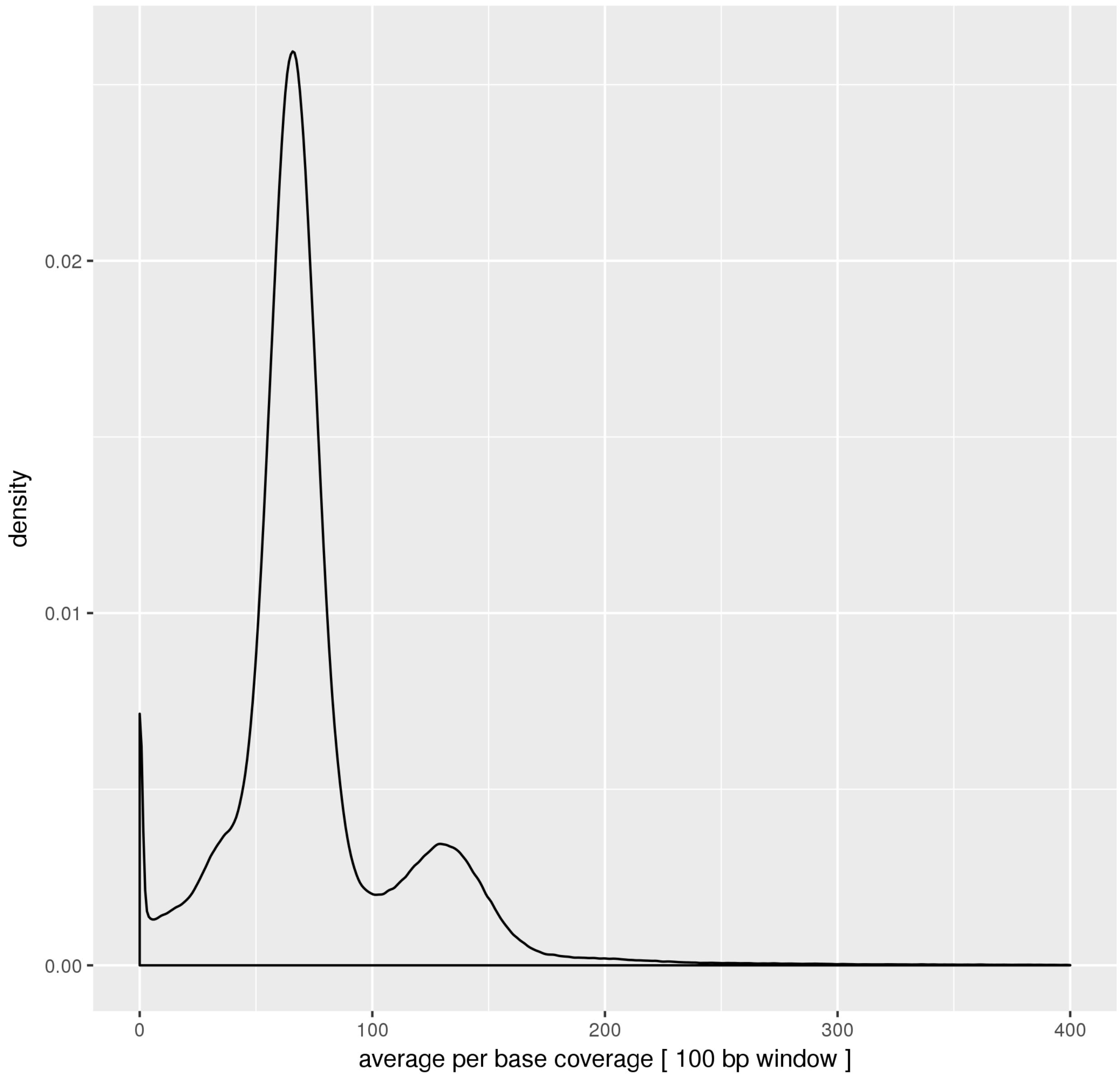


Fig. S2. Distribution of average per base coverage in 100bp windows on the *M. incognita* reference genome.

Illumina reads used to produce the reference *M. incognita* genome assembly [1] were mapped back to the reference genome using the same methodology as for the 11 isolates. Average per base coverage for each 100bp window was counted and the density at each coverage was plotted.

### Text S1. Test for association between SNP-based clusters and biological traits

We tested for significant association between the SNP-based clusters of the different *M. incognita* isolates and the following biological traits:

- Pattern of host compatibility (= host races)
- Nature of the crop plant of origin at the moment of isolation
- Geographical distribution

For the two first above-mentioned biological traits we used a Fisher's exact test while for geographical distance, we used Isolation By Distance (IBD) analysis. All these analyses were performed based on the data gathered in the Table S4 (below).

| Name | Cluster | Race | Host | Country | Region | City | GPS coordinates |
|---|---|---|---|---|---|---|---|
| R1-2 | A | 1 | Soybean | Brazil | PR | Londrina | 23°18'22.0"S 51°10'07.9"W |
| R3-2 | B | 3 | Soybean | Brazil | PR | Londrina | 23°18'22.0"S 51°10'07.9"W |
| R4-4 | B | 4 | Cotton | Brazil | SP | Vargem Grande do Sul | 21°49'52.9"S 46°53'24.3"W |
| R2-1 | C | 2 | Tobacco | Brazil | SC | Sombrio | 29°05'54.0"S 49°37'50.5"W |
| R2-6 | C | 2 | Coffee | Brazil | PR | São Jorge do Patrocínio | 23°43'47.2"S 53°56'38.4"W |
| R1-3 | C | 1 | Cucumber | Brazil | SP | Piracicaba | 22°44'05.1"S 47°38'48.2"W |
| R1-6 | C | 1 | Tobacco | Brazil | PR | Mercedes | 24°27'13.1"S 54°10'04.5"W |
| R4-3 | C | 4 | Watermelon | Brazil | PR | Londrina | 23°18'22.0"S 51°10'07.9"W |
| R3-1 | C | 3 | Cotton | Brazil | PR | Umuarama | 23°46'22.0"S 53°18'34.4"W |
| R4-1 | C | 4 | Cotton | Brazil | MT | Campo Verde | 15°33'32.0"S 55°10'04.3"W |
| R3-4 | C | 3 | Cotton | Brazil | PR, MS, BA | Umuarama, Londrina, Dourados, L.E. Magalhães | NA: pool of species |
| A14 | A | NA | Tomato | Libya | North East | NA | 32°43'51.6"N 22°07'47.4"E |
| L9 | A | NA | NA | Ivory Coast | NA | NA | 7°42'42.4"N 5°02'17.4"W |
| L19 | A | NA | NA | French West Indies | Guadeloupe | NA | 16°12'26.0"N 61°40'13.1"W |
| Morelos | C | 3 | NA | Mexico | Morelos | Morelos | 18°45'11.1"N 99°06'29.7"W |
| L27 | C | 1 | NA | USA | NA | NA | NA: US state not known |
| W1 | C | NA | Tomato | USA | California | Yolo County | 38°44'22.1"N 121°48'35.4"W |
| 557R | C | NA | Tomato | USA | North Carolina | NA | 35°49'30.3"N 78°39'11.9"W |
| HarC | C | NA | Grape | USA | California | NA | 34°13'03.2"N 117°32'54.8"W |
| VW6 | C | NA | Cotton | USA | California | NA | 34°13'03.2"N 117°32'54.8"W |

Table S3. Cluster assignment, host race status, host of origin and geographical origin of the 11 isolates.

### Association between genetic clusters and host races

The Fisher's exact test checks for independency between the tested variables, based on the co-occurrence counts.

We used the following contingency table of the 13 isolates (including the L27 isolate characterized as Race 1 and the Morelos isolate characterized as a Race 3) in clusters and host races:

| Cluster / Host Race | R1 | R2 | R3 | R4 |
|---|---|---|---|---|
| A | 1 | 0 | 0 | 0 |
| B | 0 | 0 | 1 | 1 |
| C | 3 | 2 | 3 | 2 |

The Fisher exact test returns a p-value of 1 indicating no significant association between host races and SNP clusters of isolates.

To evaluate the sensitivity of the Fisher's exact test, we also ran the same test on an artificially modified contingency table. In this modified contingency table, we assigned all the R3 to cluster B and all the R4 to cluster C and left the rest of the table unchanged.

| Cluster / Host Race | R1 | R2 | R3 | R4 |
|---|---|---|---|---|
| A | 1 | 0 | 0 | 0 |
| B | 0 | 0 | 4 | 0 |
| C | 3 | 2 | 0 | 3 |

This modification alone was enough to obtain a p-value of 0.007, indicating that this method is sensitive in detecting significant association between genetic clusters and biological traits.

### Association between genetic clusters and the nature of the crop of origin

For crop species, we performed two analyses.

- 1. All the isolates for which a crop species of origin was documented regardless the number of occurrences of the crop species (16 isolates on 8 different crops cf. Table S4).

| Cluster | Coffee | Cotton | Cucumber | Grape | Soybean | Tobacco | Tomato | Watermel |
|---|---|---|---|---|---|---|---|---|
| A | 0 | 0 | 0 | 0 | 1 | 0 | 1 | 0 |
| B | 0 | 1 | 0 | 0 | 1 | 0 | 0 | 0 |
| C | 1 | 4 | 1 | 1 | 0 | 2 | 2 | 1 |

The Fisher exact test returns a p-value of 0.69, indicating no significant association between crop species and clusters

- 2. All the isolate / crop species associations where host species count was >1 to eliminate potential biases associated to single isolate crop species (12 isolates on 4 different crops c.f. Table S4)

| Cluster | Cotton | Soybean | Tobacco | Tomato |
|---|---|---|---|---|
| A | 0 | 1 | 0 | 1 |
| B | 1 | 1 | 0 | 0 |
| C | 4 | 0 | 2 | 2 |

The Fisher exact test returns a p-value of 0.26, indicating, here again, no significant association between crop species and clusters.

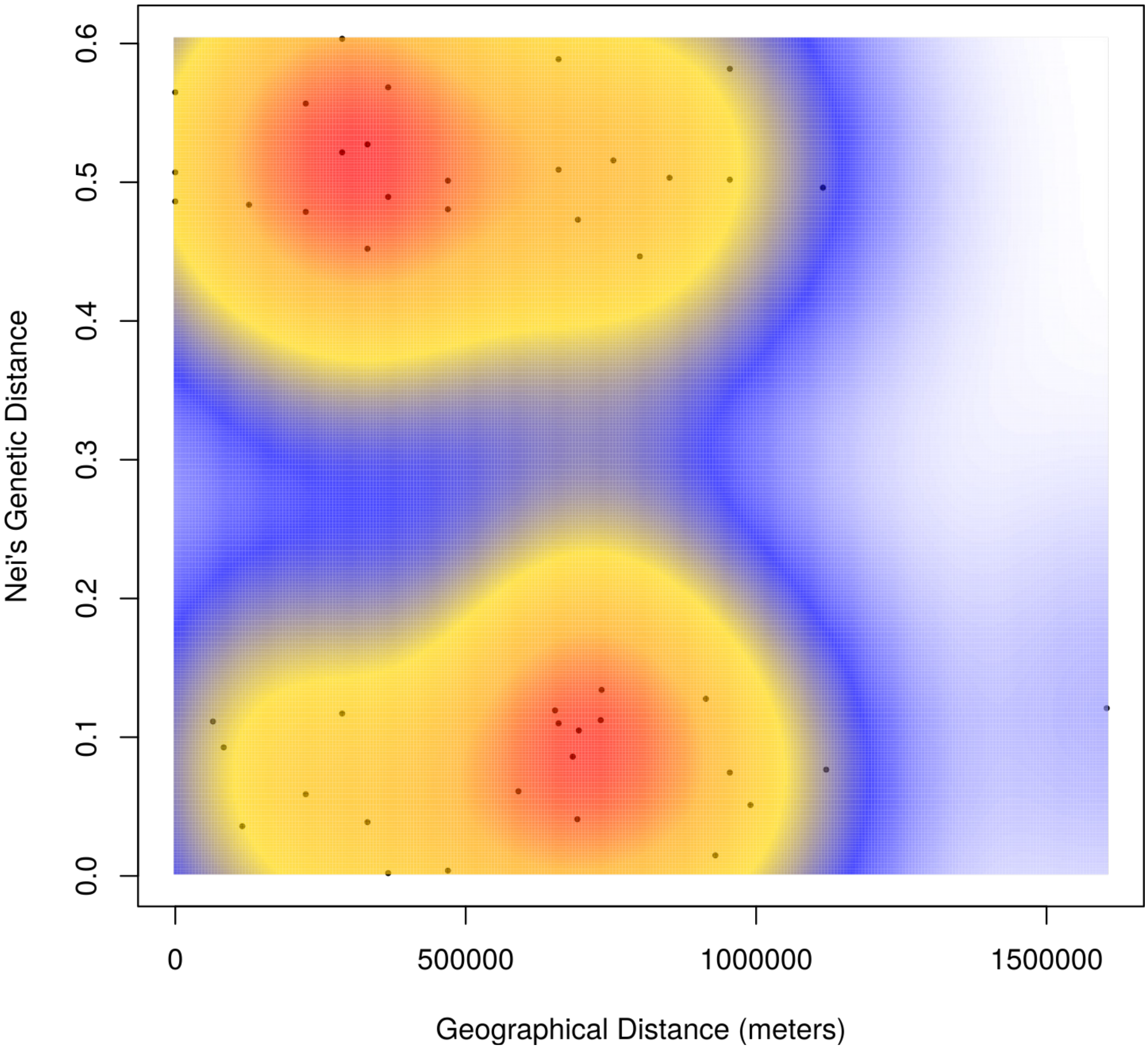


Fig. S3. Isolation by Distance (IBD) analysis between the 11 Brazilian isolates.

Each pair of isolates is plotted in a scatterplot were the x-axis is the geographical distance and the y-axis the genetic distance. Density of points is represented by different colours (blue - low, red - high).

| | Cluster A | Cluster B | Cluster C |
|---|---|---|---|
| Cluster A | X | 0.98088 | 0.3523 |
| Cluster B | 0.9904 | X | 0.52805 |
| Cluster C | 0.83675 | 0.84255 | X |

Table S4. Fixation index ($F_{ST}$) based on homozygous SNPs (upper part mean values, lower part weighted values).

### Fig. S5. Phylogenetic analysis based on SNPS present in the 14 biggest scaffolds with enough variable positions in coding regions

Unrooted topologies of all the ML trees constructed from SNPs present in coding regions for the 14 longest scaffolds containing such SNPs in enough abundance.

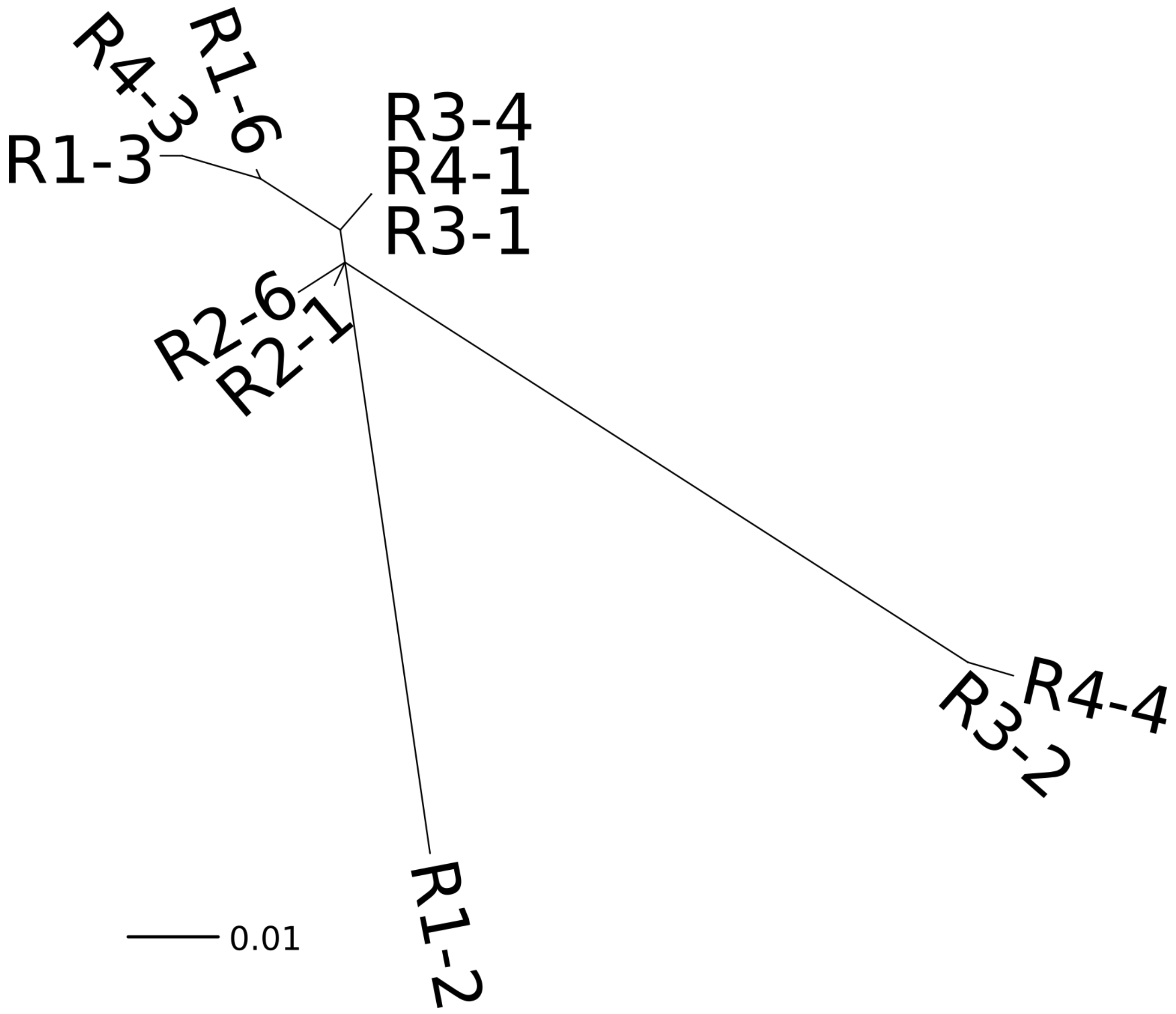


Fig. S6. Phylogenetic analysis based on SNPs in the mitochondrial genome

The tree topology exactly recapituclates the one obtained at the nuclear level.

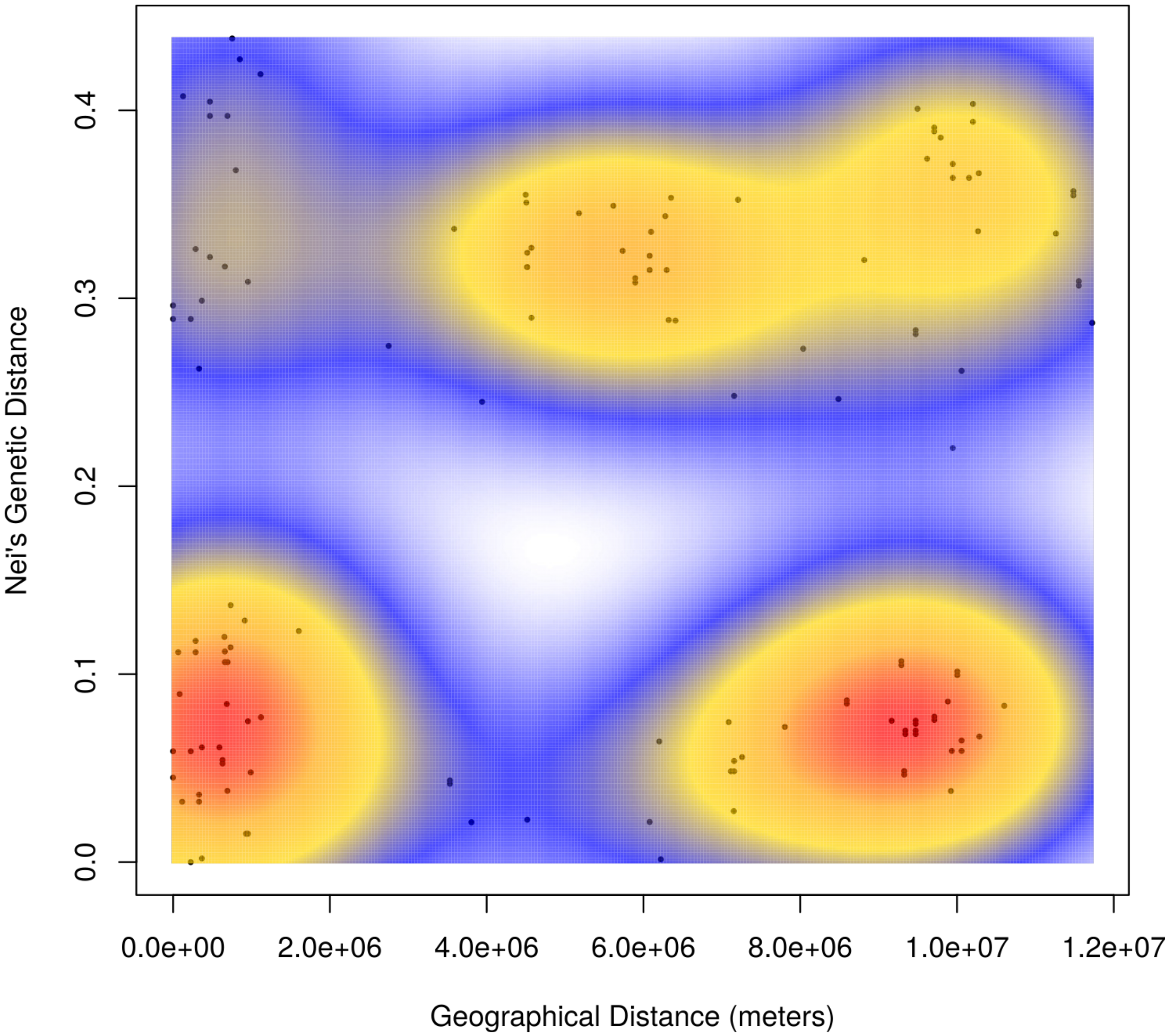


Fig. S7. Isolation by Distance (IBD) analysis between all the *M. incognita* isolates available worldwide.

Each pair of isolates is plotted in a scatterplot were the x-axis is the geographical distance and the y-axis the genetic distance. Density of points is represented by different colours (blue - low, red - high).

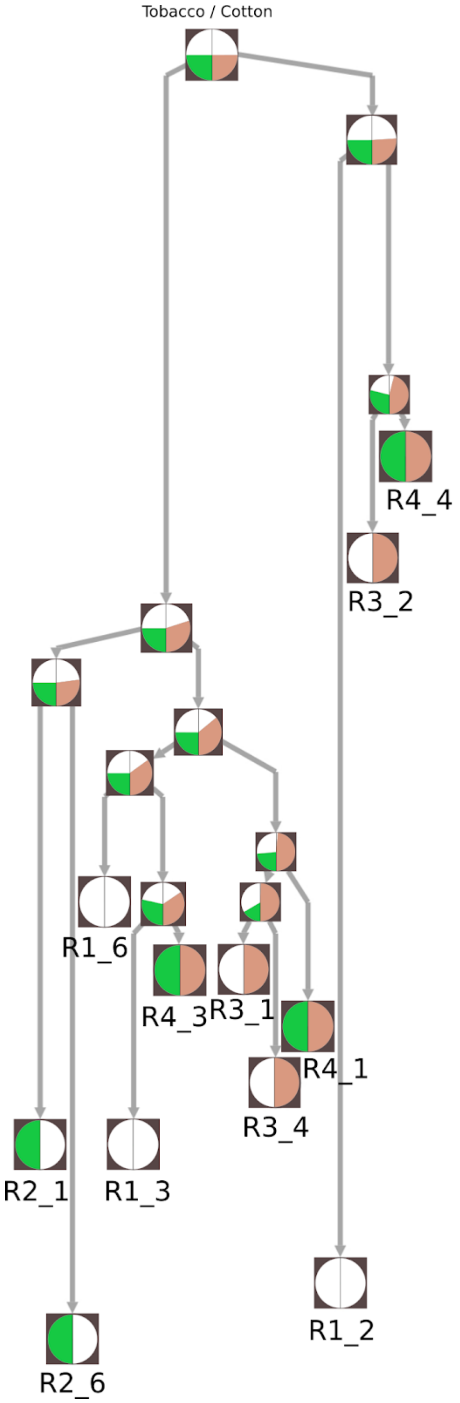


Fig. S8. Reconstruction of the ancestral states using PastML with the ML MPAA method.

*M. incognita* host races (R1-R4) are defined by differential compatibilities on Tobacco and Cotton. The left semicircle displays the probability of node being able to parasitise tobacco while the right semicircle displays the probability of the node being able to parasitise cotton.

| **Species** | **Primer** | **Sequence (5’→3’)** | **Length (bp)** | **Reference** |
|---|---|---|---|---|
| ***M. exigua*** | ex-D15F | CATCCGTGCTGTAGCTGCGAG | 562 | Randig et al., 2002 |
| | ex-D15R | CTCCGTGGGAAGAAAGACTG | | |
| ***M. incognita*** | inc-K15F | GGGATGTGTAAATGCTCCTG | 399 | Randig et al., 2002 |
| | inc-K15R | CCCGCTACACCCTCAACTTC | | |
| ***M. paranaensis*** | par-C09F | GCCCGACTCCATTTGACGGA | 208 | Randig et al., 2002 |
| | par-C09R | CCGTCCAGATCCATCGAAGTC | | |
| ***M. javanica*** | Fjav | GGTGCGCGATTGAACTGAGC | 670 | Zijlstra et al., 2000 |
| | Rjav | CAGGCCCTTCAGTGGAACTATAC | | |

Table S5. List of the SCAR molecular markers used for *Meloidogyne* spp. confirmation
