## Supplementary figures and images for "Population genomics supports clonal reproduction and multiple gains and losses of parasitic abilities in the most devastating nematode plant pest"

### Figure S5

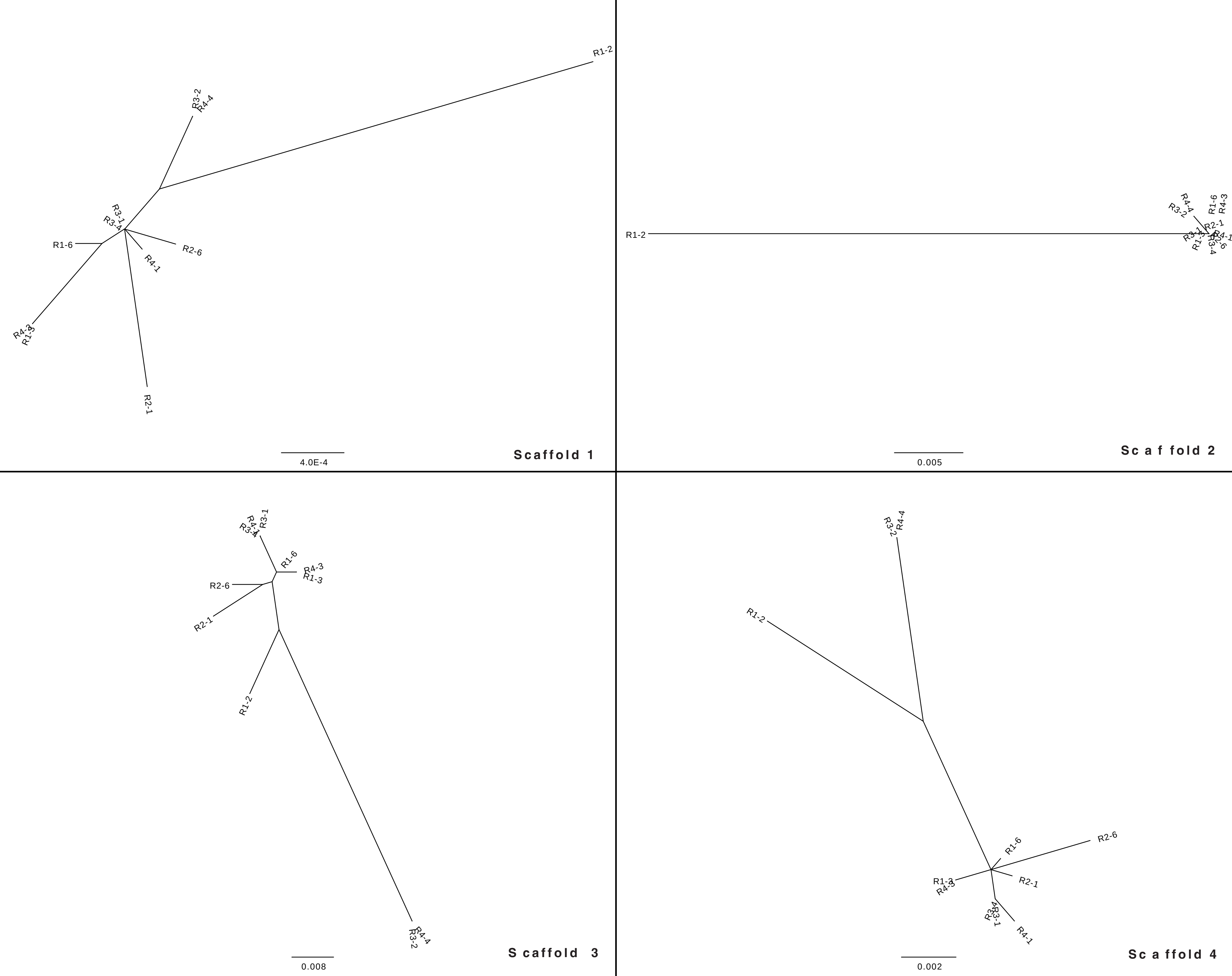

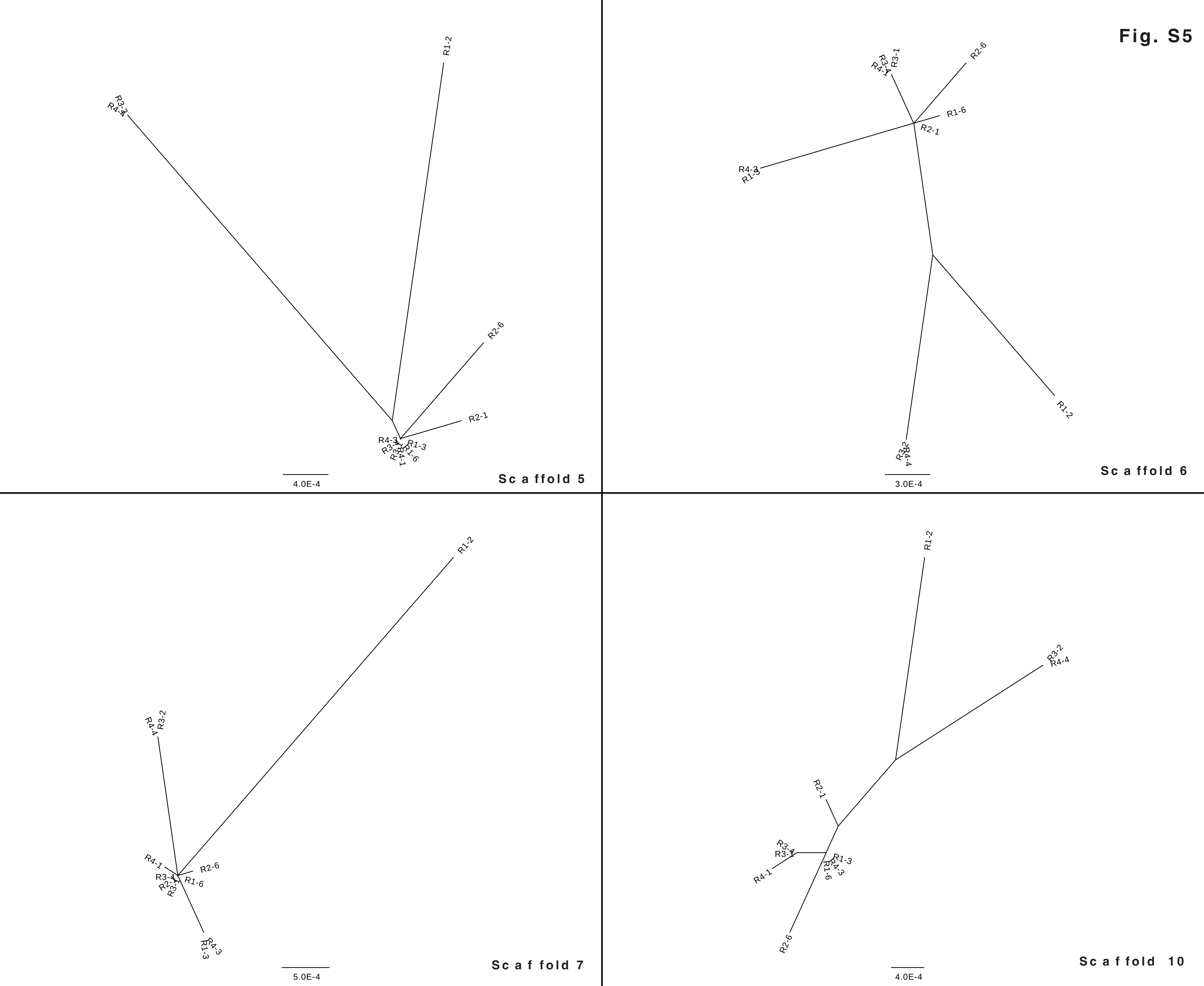

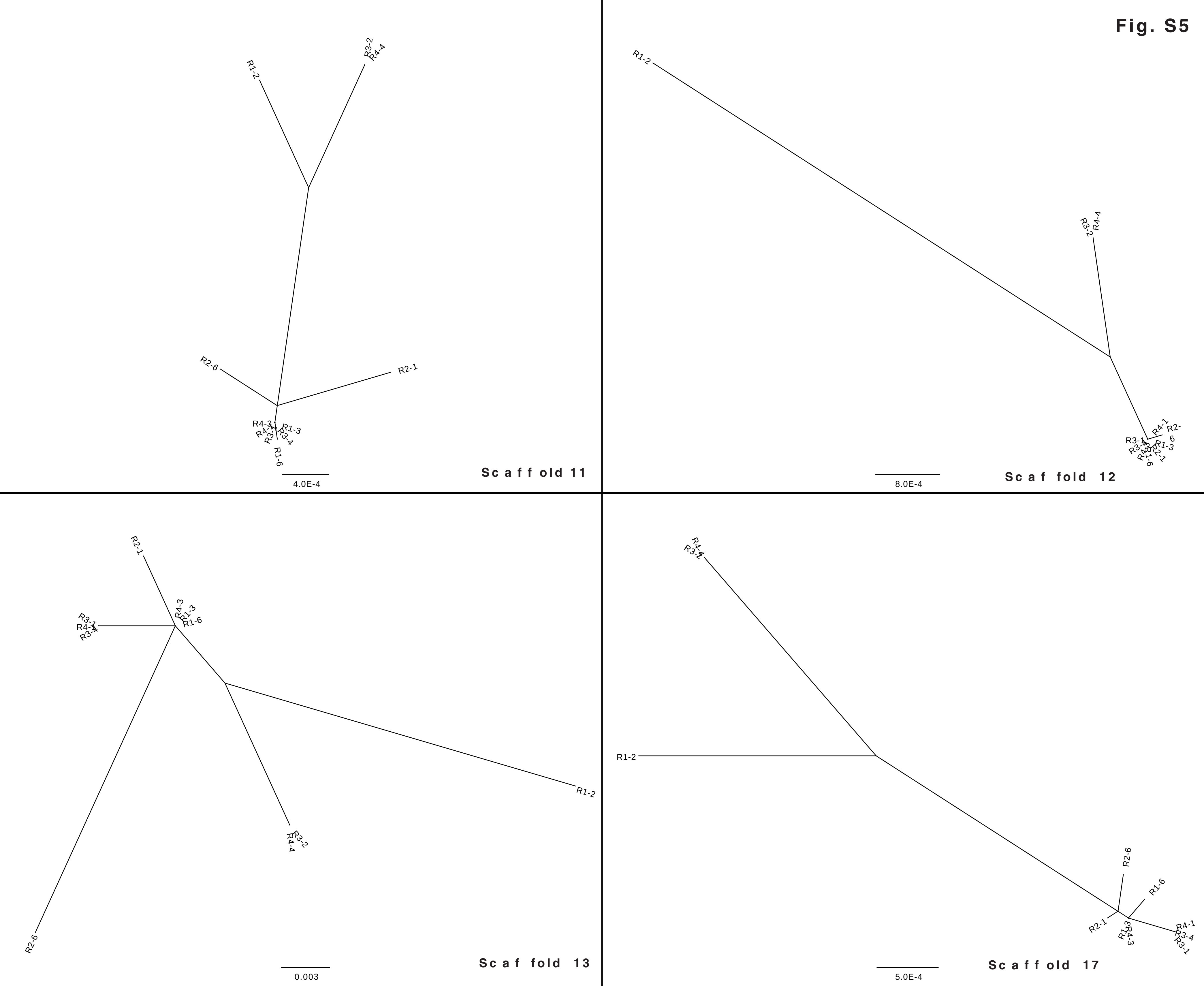

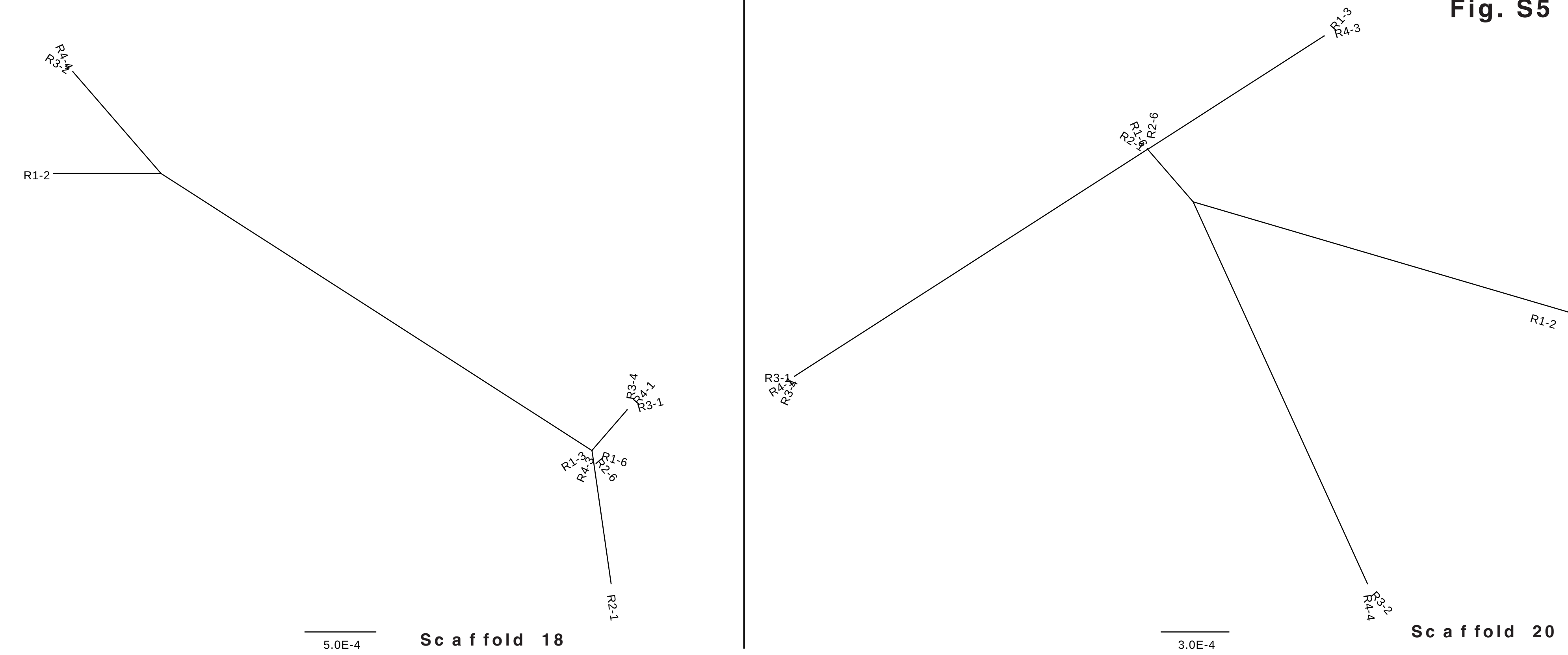
